# Coexistence of asynchronous and clustered dynamics in noisy inhibitory neural networks

**DOI:** 10.1101/2024.02.13.580163

**Authors:** Yannick Feld, Alexander K. Hartmann, Alessandro Torcini

## Abstract

A regime of coexistence of asynchronous and clustered dynamics is analyzed for globally coupled homogeneous and heterogeneous inhibitory networks of quadratic integrate-and-fire (QIF) neurons subject to Gaussian noise. The analysis is based on accurate extensive simulations and complemented by a mean-field description in terms of low-dimensional *next generation* neural mass models for heterogeneously distributed synaptic couplings. The asynchronous regime is observable at low noise and becomes unstable via a sub-critical Hopf bifurcation at sufficiently large noise. This gives rise to a coexistence region between the asynchronous and the clustered regime. The clustered phase is characterized by population bursts in the *γ*-range (30-120 Hz), where neurons are split in two equally populated clusters firing in alternation. This clustering behaviour is quite peculiar: despite the global activity being essentially periodic, single neurons display switching between the two clusters due to heterogeneity and/or noise.

## 1. Introduction

Since the pioneering studies of Winfree synchronization phenomena in biological populations are usually addressed in the context of coupled oscillators [1]: synchronization is associated to the emergence of an unique group of oscillators displaying a coherent dynamics [2]. Besides this phenomenology, one can observe also *clustering* phenomena, where the population breaks down in groups of elements displaying some sort of coherent evolution [3].

A paradigmatic complex system where these phenomena have been largely investigated are brain circuits. In this framework synchronization among a group of neurons can induce the emergences of collective oscillations (COs) [4, 5]. In this context, the existence of inhibitory interactions is fundamental in order to promote fast collective oscillatory behaviours in several areas of the brain, in particular in the hippocampus and the neocortex [6, 7].

In real systems, and in particular in brain circuits, noise is unavoidable, therefore many analyses have been devoted to its influence on coherent dynamics. In particular, a common noise source can induce synchronization and clustering phenomena, as shown for globally coupled or even uncoupled limit-cycle oscillators [8, 9, 10, 11]. This peculiar synchronization induced by common noise is referred in neurophysiology as “reliability” [12].

But also for the case of independent noise, clustering phenonema have been reported for heterogenous excitable systems with random coupling strenghts for sufficiently broad distributions of the couplings [13]. Furthermore, clustering instabilities have been shown to affect the synchronized regime in homogeneous inhibitory networks of spiking neurons subject to additive noise [5].

In this paper, we analyze in details the role of noise in promoting a regime of coexistence among clustered and asynchronous dynamics in spiking neural networks. This is a particularly relevant regime, since brain dynamics in the awake state is typically characterized by an asynchronous activity of the neurons. However, oscillations in the *γ*-range (30-120 Hz) can occasionally emerge in relation with information processing, behaviour and learning [14, 15, 16, 17]. In particular, we consider an inhibitory spiking neural network of quadratic integrate-and-fire (QIF) neurons [18] subject to Gaussian noise. The QIF model is quite general, since it represents the normal form describing the dynamics of all class I neurons in proximity to a saddle-node on a limit cycle bifurcation [19]. Furthermore, for heterogenous QIF networks exact low-dimensional mean-field (neural mass) models can be derived in terms of experimentally measurable quantities such as the population firing rate and the average membrane potential [20]. Recently, this approach has been extended to encompass extrinsic and endogenous sources of fluctuations (noise) leading to a hierarchy of low-dimensional neural mass models [21]. For his innovation with respect to classical neural mass models (e.g. the Wilson-Cowan one) this class of mean-field models has been termed *next generation* neural mass models (for the many possible applications in neural dynamics see [22]).

We will combine this mean-field analysis with accurate numerical simulations [23] to characterize at a macroscopic and microscopic level the coexisting dynamical regimes, as well as the stability of the asynchronous regime and the bifurcations associated to the emergence of the clustered state. To be more specific, the paper is organised as follows. Section 2 is devoted to the introduction of the QIF network model and the corresponding mean-field reduction methodology. The macroscopic and microscopic indicators employed to characterize the coherence and regularity of the neuronal dynamics are presented in Subsection 2.3 together with a new stability criterion for the asynchronous state in finite networks inspired by the basin stability analysis [24]. The linear stability of the neural mass models is analytically evaluated in Section 3 for Gaussian and Lorentzian noise. The asynchronous and clustered dynamics are examined in details for heterogenous synaptic couplings in Subsection 4.1 by combining accurate network simulations and neural mass results. The investigation is extended in Subsection 4.2 to homogenous couplings but relying only on network simulations. Spectral analysis of the collective oscillations is reported in Subsection 4.3 and a brief summary of the obtained results can be found in Section 5. Finally, Appendix A reports details on the numerical simulations, while Appendix B is devoted to the introduction of a criterion to select the optimal integration time step in noisy systems.

## 2. Models and indicators

### 2.1. Network Model

We consider an inhibitory population of *N* globally coupled QIF neurons [25], whose membrane potential evolution is described by the following set of equations

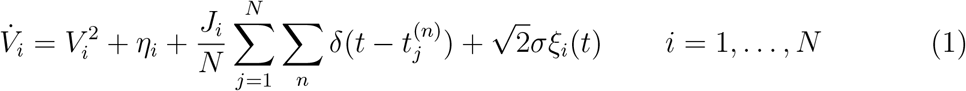

where *V*_*i*_ is the membrane potential of the *i*-th neuron, *J*_*i*_ *<* 0 the inhibitory synaptic coupling strenght and *η*_*i*_ the neuronal excitability. Whenever a membrane potential *V*_*j*_ reaches infinity a spike is emitted and *V*_*j*_ is reset to −∞. The *n*-th spike-time of neuron *j* is denoted by 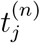.

Each neuron is subject to a common synaptic current *J*_*i*_*s*(*t*), where

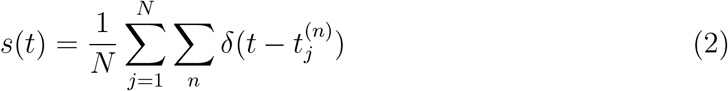

represents the activity of the network, as well as to an independent noise term of amplitude 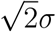, where *ξ*_*i*_(*t*) is a random Gaussian variable with ⟨*ξ*_*i*_(*t*)*ξ*_*j*_(0)⟩ = *δ*_*ij*_*δ*(*t*).

In the absence of synaptic couplings and of external noise, a QIF neuron displays excitable dynamics for *η*_*i*_ *<* 0, while it behaves as an oscillator with period 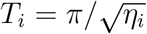 for positive *η*_*i*_. For sake of simplicity we will assume homogenous excitabilities, by fixing *η*_*i*_ = *η*_0_ = 4.2, thus all the uncoupled neurons will be supra-threshold.

In the following we will consider either heterogeneous quenched random couplings following a Lorentzian distribution (LD)

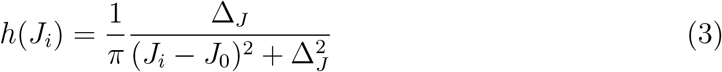

or homogeneous couplings *J*_*i*_ ≡ *J*_0_.

In order to characterize the macroscopic behaviour of the network two indicators will be essential to allow for a comparison with the mean-field formulation reported in the next Subsection. One is the mean network activity *s*(*t*) (2) and the other the mean membrane potential, defined as follows

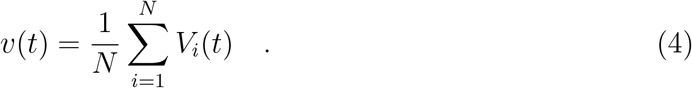

The identification of the spike-times is subject to some finite thresholding and the numerical integration of the set of stochastic differential equations Eq. (1) is explained in details in Appendix A. The model is dimensionless, however to report the times (frequencies) in physical units, we will assume as a timescale for our dynamics *τ*_*m*_ = 10 ms, corresponding to the membrane time constant.

### 2.2. Next Generation Neural Mass Model

In the recent years, it has been shown that an exact low-dimensional mean-field formulation can be developed for fully coupled networks of heterogeneous QIF neurons, with Lorentzian distributed heterogeneities [26, 27, 20]. This mean-field (neural mass) model describes the macroscopic dynamics of the network in the limit *N* → ∞ in terms of the mean membrane potential *v* (4) and the population firing rate *r*, which corresponds to the network activity *s*(*t*) (2). The main assumption of this approach is that the distribution of the membrane potentials is also Lorentzian at any time [20].

This *Lorentzian Ansatz* (LA) is violated if the neurons are randomly connected and/or in presence of noise. These more general cases can be treated by introducing a hierarchy of neural mass models taking in account the distortions to the LD of the membrane potentials [21]. Here we will briefly report the main steps to derive such mean-field formulation in the case of fully coupled inhibitory network of QIF neurons subject to additive noise.

In full generality, we can assume that both parameters *η*_*i*_ (*J*_*i*_) are distributed according to a LD *g*(*η*) (*h*(*J*)) with median *η*_0_ (*J*_0_) and half width at half maximum (HWHM) Δ_*η*_ (Δ_*J*_). In the thermodynamic limit, the network dynamics Eq. (1) can be characterized in terms of the probability density function (PDF) *p*(*V, t*|***x***) with ***x*** = (*η, J*), which obeys the following Fokker–Planck equation (FPE):

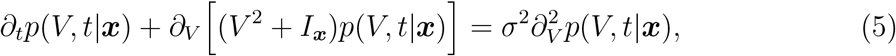

where *I*_***x***_ ≡ *η* + *Jr*(*t*). In Ref. [20] the authors assumed that in the absence of noise the solution of Eq. (5) converges to a LD for any initial PDF *p*(*V*, 0|***x***), i.e., it becomes

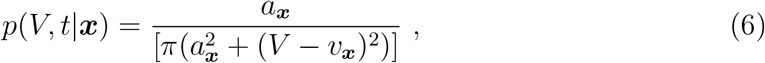

where *v*_***x***_ is the mean membrane potential and

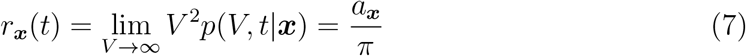

is the firing rate for the ***x***-subpopulation. The above LA joined with the assumption that the parameters *η* and *J* are independent and Lorentzian distributed lead to the derivation of exact low-dimensional neural mass models for globally coupled deterministic QIF networks [20].

Following what was done in [20] and extending it to noisy systems [21], one can introduce the characteristic function for *V*_***x***_, i.e. the Fourier transform of its PDF, namely

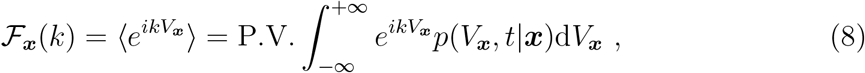

where P.V. indicates the Cauchy principal value. In this framework the FPE Eq. (5) can be rewritten as

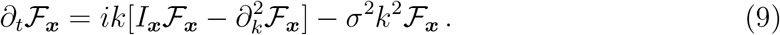

Under the assumption that ℱ_***x***_(*k, t*) is an analytic function of the parameters ***x*** one can estimate the characteristic function averaged over the heterogenous population

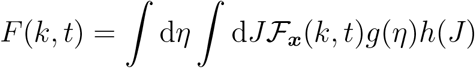

and via the residue theorem the corresponding FPE, namely

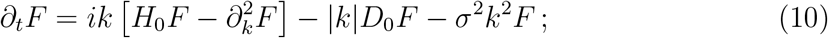

where *H*_0_ = *η*_0_ + *J*_0_*r* and *D*_0_ = Δ_*η*_ + Δ_*J*_ *r*.

For the logarithm of the characteristic function, Φ(*k, t*) = ln(*F* (*k, t*)), one obtains the evolution equation

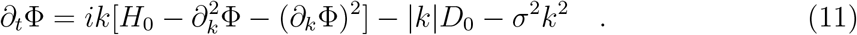

In this context the LA amounts to Φ_*L*_ = *ikv* − *a*|*k*|. By substituting Φ_*L*_ in Eq. (11) for *σ* = 0 one gets

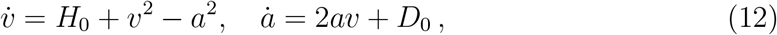

which coincides with the two dimensional exact MF reported in [20] with *r* = *a/π*.

In order to consider deviations from the LD, the authors of Ref. [21] analysed the following general polynomial form for Φ

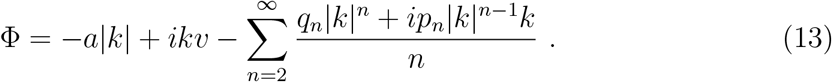

and introduced the notion of complex *pseudo-cumulants*, defined as follows

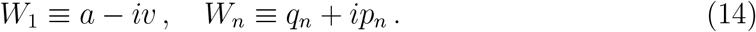

By inserting the expansion Eq. (13) in the Eq. (11) one gets the evolution equations for the pseudo-cumulants, namely:

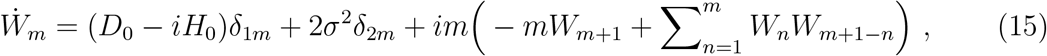

where *δ*_*ij*_ is the Kronecker delta function and for simplicity we assumed *k >* 0. It is important to notice that the time-evolution of *W*_*m*_ depends only on the pseudo-cumulants up to the order *m* + 1, therefore the hierarchy of equations can be easily truncated at the *m*-th order by setting *W*_*m*+1_ = 0. As shown in Ref. [21] the modulus of the pseudo-cumulants scales as |*W*_*m*_| ∝ *σ*^2(*m*−1)^ with the noise amplitude, therefore it is justified to consider an expansion limited to the first few pseudo-cumulants.

In this paper, we will consider (15) up to the third order to obtain the corresponding neural mass model, i.e.

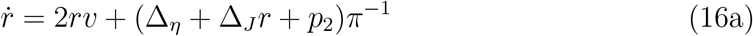

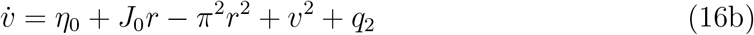

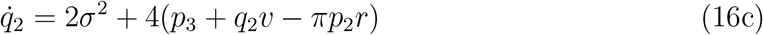

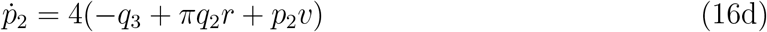

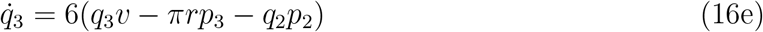

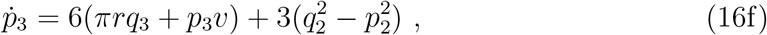

with the closure *p*_4_ = *q*_4_ = 0. The macroscopic variables *r* and *v* represent the population firing rate and the mean membrane potential, while the terms *q*_2_, *p*_2_, *q*_3_, *p*_3_ take in account the dynamical modification of the PDF of the membrane potentials with respect to a Lorentzian profile. Besides the third-order neural mass models, we will also consider the second-order one, which can simply be obtained by considering Eqs. (16a,16b,16c,16d) by setting *q*_3_ = *p*_3_ ≡ 0

Since Eq. (16) is a set of deterministic ordinary differential equations, one can use standard numerical methods to integrate them. In particular, we employed a 4th order Runge-Kutta method [28]. The neural mass results will be compared with network simulations in the following and employed to initialize the network in an asynchronous state (see, e.g., Section 2.3.2 and Appendix B).

### 2.3. Indicators

#### 2.3.1. Macroscopic Indicators

The evolution of the membrane potential of a neuron, in particular in the supra-threshold regime, can be interpreted as the rotation of the phase of an oscillator and many models have been derived by employing such an analogy. In terms of these phases the level of synchronization of the oscillators (neurons) can be measured in terms of macroscopic order parameters that we will introduce in the following.

For the QIF model, the membrane potential *V*_*i*_ of the *i*-th neuron is usually mapped in the phase *θ*_*i*_ of an oscillator via the following transformation

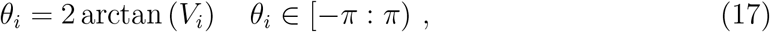

that leads from the QIF network Eq. (1) to the equivalent *θ*-neuron network [18]:

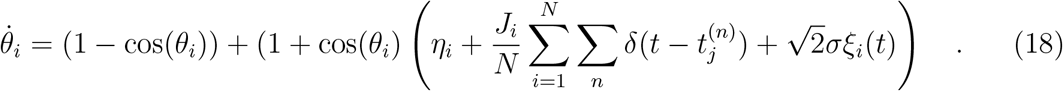

Unfortunately, the distribution of the phases *P* (*θ*_*i*_) is not uniform even for uncoupled neurons, since 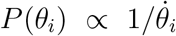. This can therefore lead to apparent synchronization phenomena in the *θ*-space in noisy enviroments [29, 30].

In order to avoid such a problem, the phase *θ*_*i*_ of the *i*-th neuron at a certain time *t* can be obtained simply by interpolating linearly between the previous and the next spike time of the considered neuron, as follows

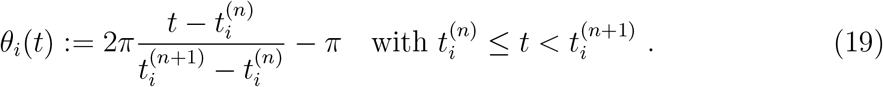

where 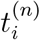 is the time at which the *n*-th spike is emitted by *i*-th neuron.

Now that the phases have been defined, we can introduce suitable order parameters to measure the level of phase synchronization in the network [31, 32, 33, 34, 35], in particular we consider the so-called *Kuramoto-Daido order parameters*

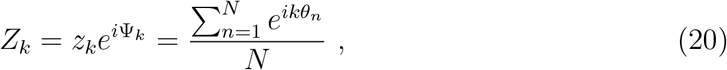

where *Z*_*k*_ is a complex number and *z*_*k*_ and Ψ_*k*_ are the corresponding modulus and phase. For *k* = 1 the usual Kuramoto order parameter [31] is recovered. For a network of *N* oscillators one expects 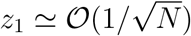 in the asynchronous regime and *z*_1_ will be finite (one) for a partially (fully) synchronized state. Unfortunately, *z*_1_ is also exactly zero when the oscillators are equally divided in 2 perfectly synchronized clusters in anti-phase. To characterize regimes presenting *k* clusters, Daido [32] introduced the parameters *Z*_*k*_. Indeed, *z*_*k*_ will be one whenever the system presents *k* clusters equally spaced in phase and equally populated, while *z*_*k*_ will approach zero for a sufficiently large network if the phases are evenly distributed over the whole interval.

To denote that the order parameters are estimated by employing the phases defined in terms of the the spiking times as in Eq. (19) we will use a super-script (*s*), while the lack of a super-script will denote the use of the phases defined as in Eq. (17).

In the mean-field framework previously introduced in Section 2.2, *Z*_*k*_ can be obtained as follows [36]

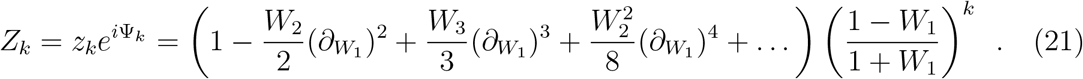

Note that the corrections obtained from the higher order pseudo cumulants *W*_*j*_, i.e. with *j >* 3, should be negligible. In absence of noise and for the usual Kuramoto order parameter *Z*_1_ this reduces to the following conformal transformation

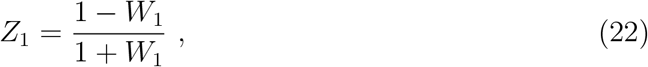

as shown in Ref. [20].

Another macroscopic indicator able to distinguish asynchronous regimes from oscillatory ones characterized by a partial synchrony of the neurons is the variance Σ_*v*_ of the mean membrane potential *v* estimated over a certain time window *T*_*W*_ . This quantity is expected to be vanishing small 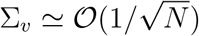 in the asynchronous regime and finite whenever COs are present.

In order to identify the asynchronous and partially synchronized regime, since these are characterized by definitely different values of Σ_*v*_, we can define a threshold value *S*_*θ*_ and whenever Σ_*v*_ *< S*_*θ*_ (Σ_*v*_ ≥ *S*_*θ*_) the dynamics will be identified as asynchronous (partially synchronized). The threshold value *S*_*θ*_ is usually chosen as the mean of the values measured in the asynchronous and partially synchronized regimes, however the identification of the regimes is quite insensitive to the exact value of *S*_*θ*_.

Also the Kuramoto-Daido order parameter can be employed for this discrimination, and as we will see in the following for the examined dynamics the most suitable indicator will be 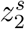.

#### 2.3.2. Stability Criterion

The considered model exhibits in a certain parameter interval a region of coexistence of two different dynamical regimes: an asynchronous and a partially synchronous one. Our goal is to quantify the stability of the asynchronous regime for the finite network by varying the noise intensity. Therefore, we have introduced the following criterion inspired by the basin stability criterion [24], which has been applied in many different contexts [37, 38, 39, 40, 41].

The main idea is to consider a solution of a system, perturb this solution several times with a magnitude that is given by a parameter and let the dynamics evolve each time. Then one measures the fraction of how often the system evolves back to a desired solution. Here, we proceed as follows: we initialize the values of the membrane potentials {*V*_*i*_} according to

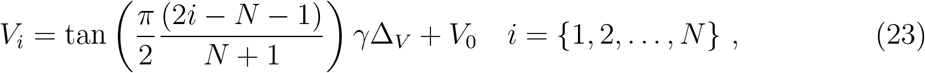

with *V*_0_ = *v*^*^ and Δ_*V*_ = *πr*^*^, where (*v*^*^, *r*^*^) are the fixed point solution of the third-order neural mass model Eq. (16). Note that for *γ* = 1 Eq. (23) will result in the Lorentzian distribution that is expected for an asynchronous regime at equilibrium, while the extreme case *γ* = 0 fixes all the membrane potentials *V*_*i*_ ≡ *V*_0_, i.e., it results in a fully synchronized initial state.

For different values of the parameter *γ* ≤ 1 we simulate the system for a time *T*_*t*_, after which we verify whether the systems is asynchronous or partially synchronized. For each value of *γ* we repeat this procedure *M* times for different noise realizations and count how many times *M*_*c*_ the system exhibits its asynchronous state after the integration time *T*_*t*_. Thus, we can measure the stability of the asynchronous state of the chosen configuration via the following indicator

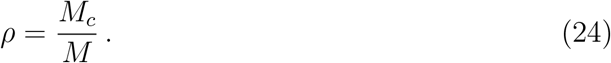

A completely stable (unstable) asynchronous regime will correspond to *ρ* = 1 (*ρ* = 0) for any value of *γ*. However, in general *ρ* will be a function of *γ*. In the bistable regime, by decreasing the *γ* value one will eventually observe a transition towards the partially synchronized regime. Thus, the value of *γ* where this “transition” happens is a measure of the stability of the bistable system.

#### 2.3.3. Characterization of irregular spiking

As we will show in the following, the partially synchronized state is characterized by two clusters of neurons firing in anti-phase. Furthermore, the neurons in each cluster do not remain in the same cluster over long time periods. Instead they tend to switch from one cluster to the other, despite the collective dynamics being always characterized by two clusters of neurons firing in alternation. Therefore the usual behavior for a neuron is to fire in correspondence with every second neuronal burst, but with irregularities in this repetition. We want to introduce a measure of these irregularities in the sequence of spikes of the neurons.

In order to develop this measure we store the sets 𝒮 = [*s*_1_ = (*t*_1_, *i*_1_), *s*_2_ = (*t*_2_, *i*_2_), *s*_3_ = …] of firing times and firing neurons in the network for a certain time interval, where *t*_*j*_ is the firing time and *i*_*j*_ ∈ [1, …, *N*], the index of the firing neuron. Moreover, we also store the bursting times *b*_*k*_ at which the neuronal bursts occur. These are identified by the maxima in the population firing rate *r*. As a convention we define the burst *k* as the collection of all the spiking events occurring within the time interval

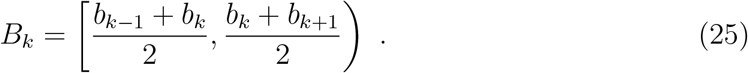

thus the spike *s*_*m*_ is emitted within the burst *k* if *t*_*m*_ ∈ *B*_*k*_.

The regular behavior for a 2 cluster state would be that a neuron, which fires within the burst *k*, would emit its next spike within the burst *k* + 2. To analyze the eventual irregularity of the individual neurons we create a ordered list 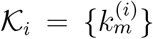 reporting the bursts within which the considered neuron *i* has fired during the observation time interval *T* . The first two bursts *k* = 0 and *k* = 1 are employed to identify if the neuron belongs to the first or second cluster, i.e. 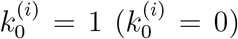 if the spike of neuron *I* occurred within *B*_1_ (*B*_0_). Then, for each neuron we introduce a counter *E*_*i*_ of “early spikes” in the following way: We go through the list 𝒦_*i*_ of bursts to which neuron *i* has contributed and we increment *E*_*i*_ by one each time 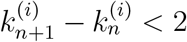. If we had a total of ℬ bursts in the considered time interval *T*, then the fraction of spikes that have been emitted too early by the neuron *i* is

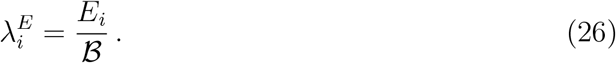

In particular if a neuron would fire during each burst we will have 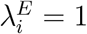.

Similarly, the fraction of “late delivered spikes” can be calculated by using a second counter *L*_*i*_ that is incremented by one each time 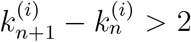, which leads to define the following fraction of late spiking neurons

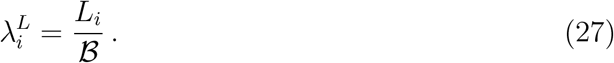

Note that 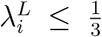 due to its definition, since the counter is incremented whenever 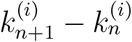 is at least 3.

In summary, the percentage of times the spikes occur outside of the expected time-frames, i.e., the percentage of irregular spikes, is

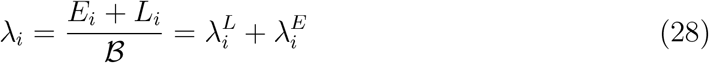

The number of neurons for which *λ*_*i*_ = 0 until the time *t* define *the surviving neurons*, i.e., those which fire regularly every 2 bursts as expected. The fraction of surviving neurons *S*(*t*) until the time *t* can be defined as

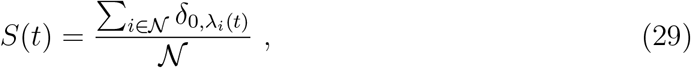

where *δ* is the Kronecker delta, and 𝒩 is the number of non-silent neurons. The silent neurons should be removed from the count, since one always has *λ*_*i*_ = 0 for those neurons: a neuron that never spikes obviously does not have any associated spike time in an unexpected time interval. For our analysis we considered *T* = 55 s.

The survival probability *S*(*t*) is usually defined as [42] :

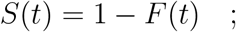

where 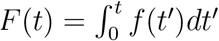 is the cumulative distribution function and *f* (*t*) the PDF of the neuronal survival times, i.e., the time until which the considered neuron fires regularly once every two bursts.

## 3. Linear stability analysis of the asynchronous state

In this Section we analyse the stability of the asynchronous regimes within the neural mass formulation. For the neural mass model Eq. (16), the asynchronous states correspond to fixed point solutions 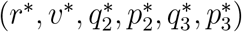. In particular, we will study the stability of these solutions by considering the linearization of Eq. (16) in proximity of the considered fixed points, namely

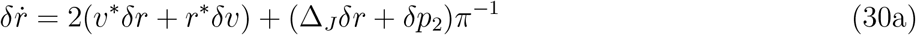

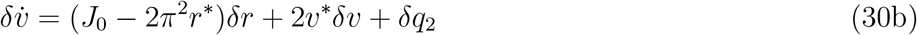

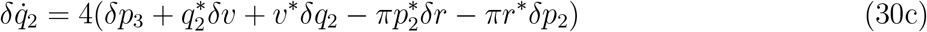

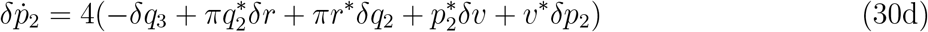

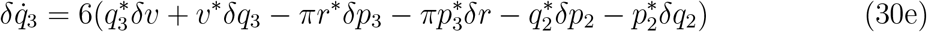

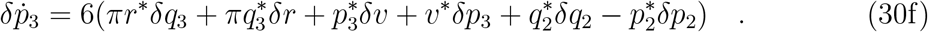

### 3.1. Deterministic Case

Let us start from the case in absence of noise. In this case the mean-field equations reduce to the exact formulation reported in [20]

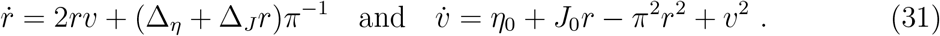

The fixed point solutions can be obtained by solving the following equations

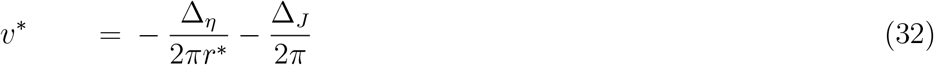

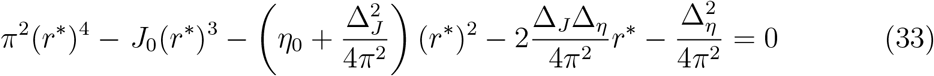

for the parameter values considered in this paper, namely inhibitory coupling *J*_0_ = −20, *η*_0_ = 4.2 and Δ_*J*_ = 0.02 and 0 ≤ Δ_*η*_ ≤ 0.30 the system exhibits 2 complex conjugate and 2 real solutions. Among the real ones only one corresponds to a positive firing rate *r*^*^ and is therefore physically acceptable.

The stability of such a solution can be obtained by analysing the linear evolution (*δr*(*t*), *δv*(*t*)) = e^*λt*^(*δr*(0), *δv*(0)) in proximity of the physical fixed point solutions (*r*^*^, *v*^*^). This amounts to solving second order characteristic equations for the eigenvalue problem associated to Eq. (30) with *p*_2_ = *q*_2_ = *p*_3_ = *q*_3_ = 0, which gives the following result

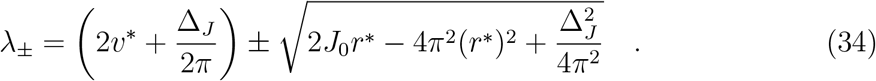

For the chosen values of the parameters the square root in Eq. (34) is always purely imaginary. Therefore the fixed point is a focus and, when inserting Eq. (32), the real part of the eigenvalues is simply given by

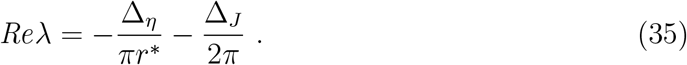

The focus is always stable, apart from the fully homogenous situation Δ_*η*_ = Δ_*J*_ = 0 in which case it becomes marginally stable. The heterogeneities tend to stabilize the focus solutions. Therefore, even in the case of homogeneous coupling Δ_*J*_ = 0, a small heterogeneity in the excitabilities measured by Δ_*η*_ is sufficient to render the fixed point stable. This can later be seen for *σ* = 0 in Fig. 2.

**Figure 1.**
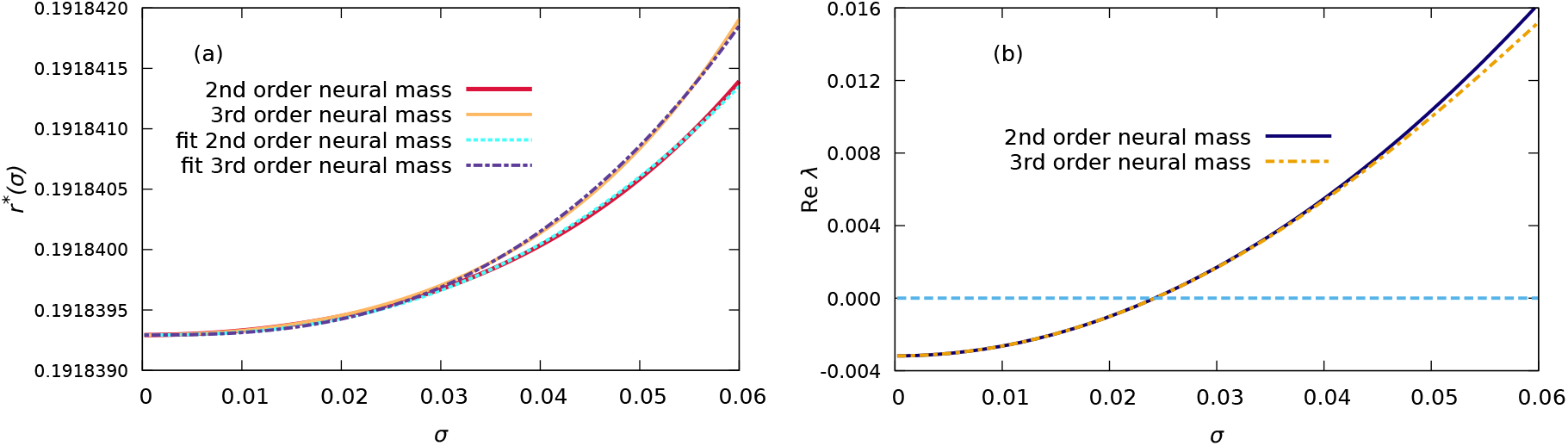
(a) Fixed point solutions for the firing rate *r*^*^ versus the noise amplitude *σ* for the 2nd and 3rd-order neural mass models. Fitting to the data with the expression *r*(*σ*) = *r*(0) + *aσ*^*α*^ are also reported. (b) Real part of the most unstable eigenvalues for the 2nd and 3rd-order neural mass model versus *σ*. Parameters are set to *η*_0_ = 4.2, *J*_0_ = −20.0, Δ_*η*_ = 0, and Δ_*J*_ = 0.02.

**Figure 2.**
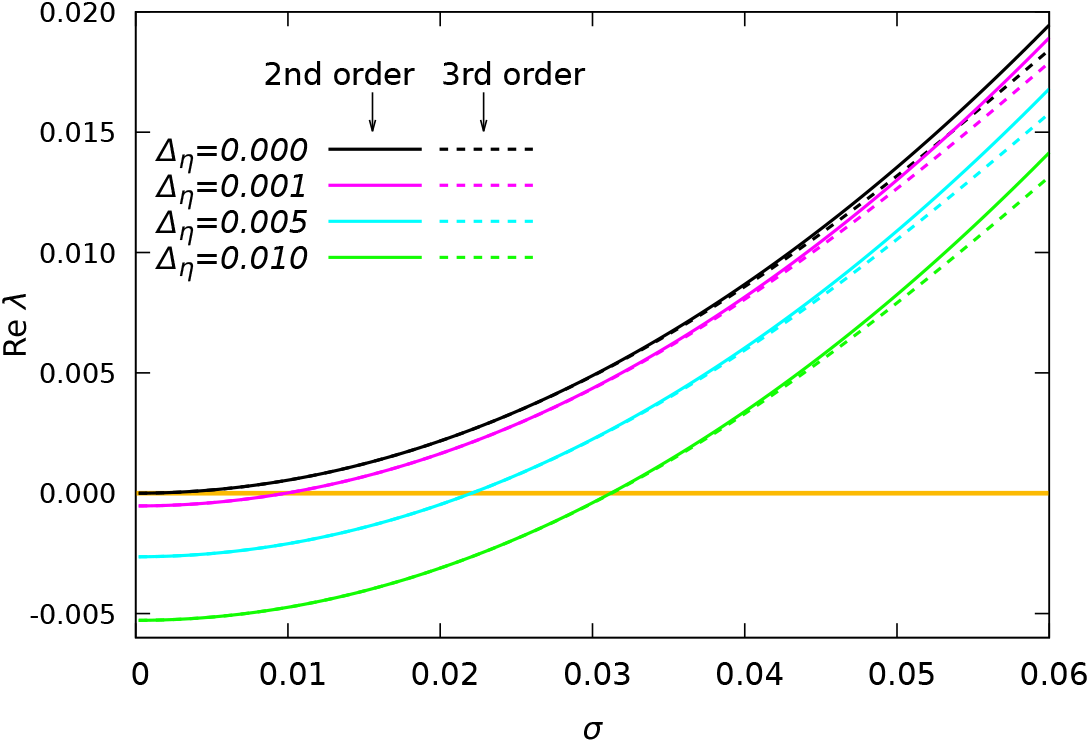
Real part of the most unstable eigenvalues for the 2nd and 3rd order neural mass model versus *σ* for different level of heterogeneity Δ_*η*_. Parameters are set to *η*_0_ = 4.2, *J*_0_ = −20.0 and Δ_*J*_ = 0.

### 3.2. Gaussian Noise

In presence of additive Gaussian noise of amplitude *σ*, we always observe a stable focus for sufficiently small *σ*. The effect of noise is an increase of the value of the firing rate *r*^*^(*σ*) with respect to the case in absence of noise *r*^*^(0). In particular, the correction to the deterministic solution can be written as *r*^*^(*σ*) ≃ *r*^*^(*σ*) + *aσ*^*α*^.

For Δ_*η*_ = 0 and the parameters usually employed in this analysis one obtains *α* ≃ 2.5 for the 2nd-order neural mass model, while the growth is even faster for the third-order model with *α* ≃ 2.7, as evident from Fig. 1 (a).

To analyze the stability of the fixed points we have identified the corresponding eigenvalues by solving the associated characteristic polynomial, that can be of the 4th (6th) order depending if we consider the neural mass model to the 2nd (3rd) order. The linear stability analysis reveals that the 4 eigenvalues for the 2nd order neural mass are two complex conjugate pairs, whose real parts are definitely negative for *σ* = 0 and Δ_*η*_ and/or Δ_*J*_ not zero, as evident from Eq. (35).

As shown in Fig. 1 (b), noise destabilizes the fixed point, since it leads to an increase of the real part of the maximal eigenvalues. In particular, these two eigenvalues can cross the zero axis at a critical noise amplitude *σ*_*H*_ . Thus indicating that the fixed point solution becomes unstable via a Hopf bifurcation giving raise to COs. For the case shown in Fig. 1 (b) we have *σ*_*H*_ ≃ 0.0243. The third order model displays 3 pairs of complex conjugates eigenvalues, however the fixed point looses stability exactly at the same *σ*_*H*_ value via a Hopf bifurcation (see Fig. 1 (b)). The effect of noise on the stability of the foci is analogous for a network with homogenous couplings (Δ_*J*_ = 0) and heterogeneous currents Δ_*η*_ *>* 0, as shown in Fig. 2. As we will see in the following these Hopf bifurcations are sub-critical, thus leading to a coexistence region between asynchronous dynamics and COs.

### 3.3. Lorentzian Noise

It is worth mentioning in this context that, by assuming that the white noise terms *ξ*_*i*_(*t*) are Lorentzian distributed, it is still possible to obtain the corresponding low-dimensional neural mass model in an exact manner [43, 44]. In particular, by assuming that the *ξ*_*i*_(*t*) random term follow a LD centered in zero and with HWHM Γ, one can obtain the following two-dimensional neural mass model [44]

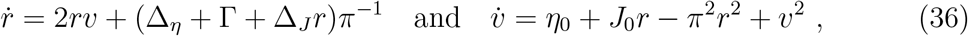

which is identical to Eq. (31) apart for the Γ term that contributes exactly as the HWHM Δ_*η*_ of the neural excitabilities {*η*_*i*_} to the mean-field dynamics. Therefore, in the thermodynamic limit the Lorentzian noise can be assimilated to a quenched disorder in the heterogeneities as shown also in Refs. [43, 44]. In the present context, this implies that the neural mass model Eq. (36) displays only stable foci solutions as Eq. (31) and no collective oscillations are observable, contrary to the case where the noise is Gaussian distributed.

## 4. Numerical results

In this Section we will analyze and characterize the clustering transition induced by noise. In particular, we will first investigate heterogenous couplings, where we fix Δ_*J*_ = 0.02, in this case we will compare the results obtained within the mean-field approach with network simulations. Successively, we will examine the homogenous situation, where Δ_*J*_ = 0, by relying only on network simulations. If not specified otherwise we fix the parameters to the following values *J*_0_ = −20, *η*_0_ = 4.2 and Δ_*η*_ = 0, and we consider an inhibitory network of size *N* = 200000 subject to Gaussian additive noise.

### 4.1. Heterogeneous synaptic couplings

In Subsection 3.2 we have shown that in the mean-field formulation the asynchrous regime remains stable up to a noise of amplitude *σ*_*H*_ ≃ 0.0243, where it destabilizes via a Hopf bifurcation. In this Subsection we will characterize such a transition and the observed regimes in full details.

#### 4.1.1. The clustering transition

In order to understand if the transition is super- or sub-critical, we perform simulations by varying quasi-adiabatically the noise amplitude *σ* (for details see Appendix A) and by measuring for each value of *σ* the variance Σ_*v*_ of the mean membrane potential.

As shown in Fig. 3, for the 2nd and 3rd order neural mass models we observe that starting from *σ* = 0 and by increasing the noise amplitude, the variance Σ_*v*_ remains zero until a value near *σ*_*H*_ is reached, as expected for a constant mean membrane potential (*v* = *v*^*^). Afterwards it jumps to some finite value due to the emergence of COs. Once the noise amplitude reaches *σ* = 0.03 the quasi-adiabatic simulations are then continued by decreasing *σ* in steps of Δ*σ*. In this case Σ_*v*_ stays finite down to values *σ*_*SN*_ ≃ 0.004 and then at even smaller value of *σ* returns to zero. This scenario is typical for a sub-critical Hopf bifurcation, characterized by the coexistence of oscillatory and stationary behaviours in the range *σ* ∈ [*σ*_*SN*_, *σ*_*H*_]. In particular, at *σ*_*SN*_ one expects that the oscillatory solution will disappear via a saddle-node bifurcation of limit cycles.

**Figure 3.**
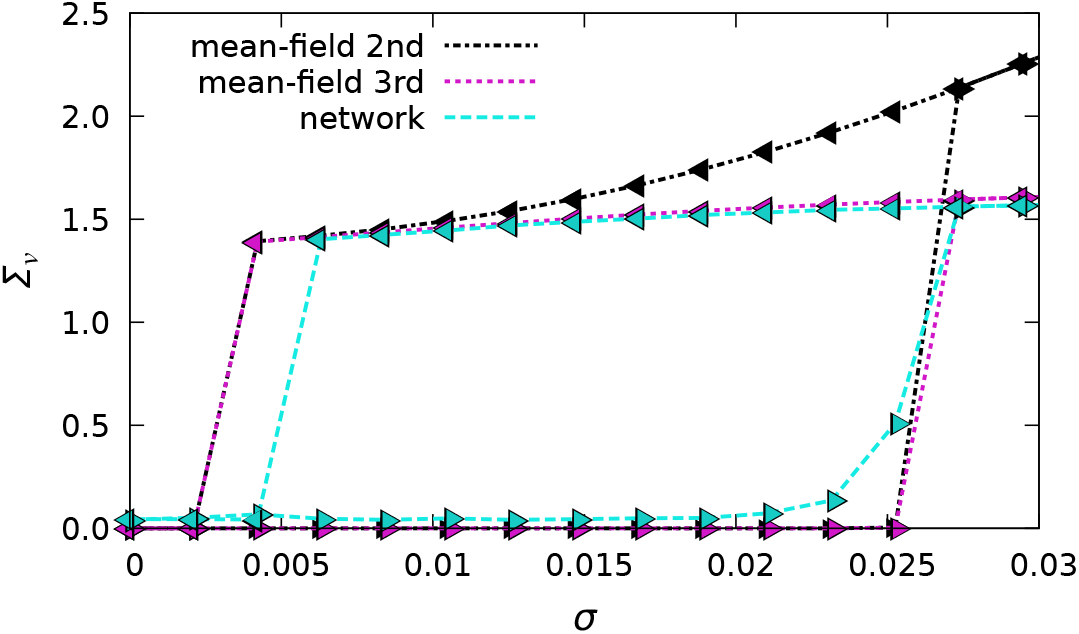
Variance Σ_*v*_ of the mean membrane potential *v* versus noise amplitude *σ* obtained via quasi-adiabatic simulations. The decrease or increase of *σ* performed during the adiabatic simulations is indicated by the direction of the triangles’ tip. The dashed lines are meant for visual aid. The parameters for the quasi-adiabatic simulations are 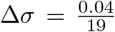, *t*_*T*_ = 20 s, *t*_*S*_ = 25 s, for the neural mass we employed *t*_*T*_ = 100 s to allow the system to better relax, and the network size was fixed to *N* = 200000. Other parameters as in Fig. 1.

The network simulations agree quite well with those of the 3rd order neural mass model, apart some finite-size effects that imply finite values 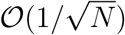 for Σ_*v*_ even in the asynchronous regime and a backward transition from the oscillatory to the asynchronous regime occurring at a larger noise amplitude, namely *σ* ≃ 0.006, instead that at *σ*_*SN*_ . In contrast, the 2nd order neural mass model displays clear differences with the 3rd order one and the network simulations in the oscillatory regime for *σ >* 0.01 (see black triangles in Fig. 3). This is probably due to an instability of the 2nd order model at large noise amplitudes.

#### 4.1.2. Coexisting regimes

Let us now examine the macroscopic properties of the asynchronous and clustered regimes in the coexistence region in more detail. In order to gain some insight we report the mean membrane potential *v* versus time for the two regimes at *σ* = 0.00842 in Fig. 4 (a and d). In the asynchronous state *v* is exactly constant for the neural mass simulations, while it displays small erratic fluctuations when obtained from network simulations. This is due to the fact that the stable fixed point is a focus, therefore the presence of finite-size fluctuations excites continuosly relaxation oscillations towards the focus. As shown in Fig. 4 (d) *v*(*t*) is periodically oscillating in the clustered regime and in this case the network simulations agree quite well with the neural mass results obtained for both 2nd and 3rd order models.

**Figure 4.**
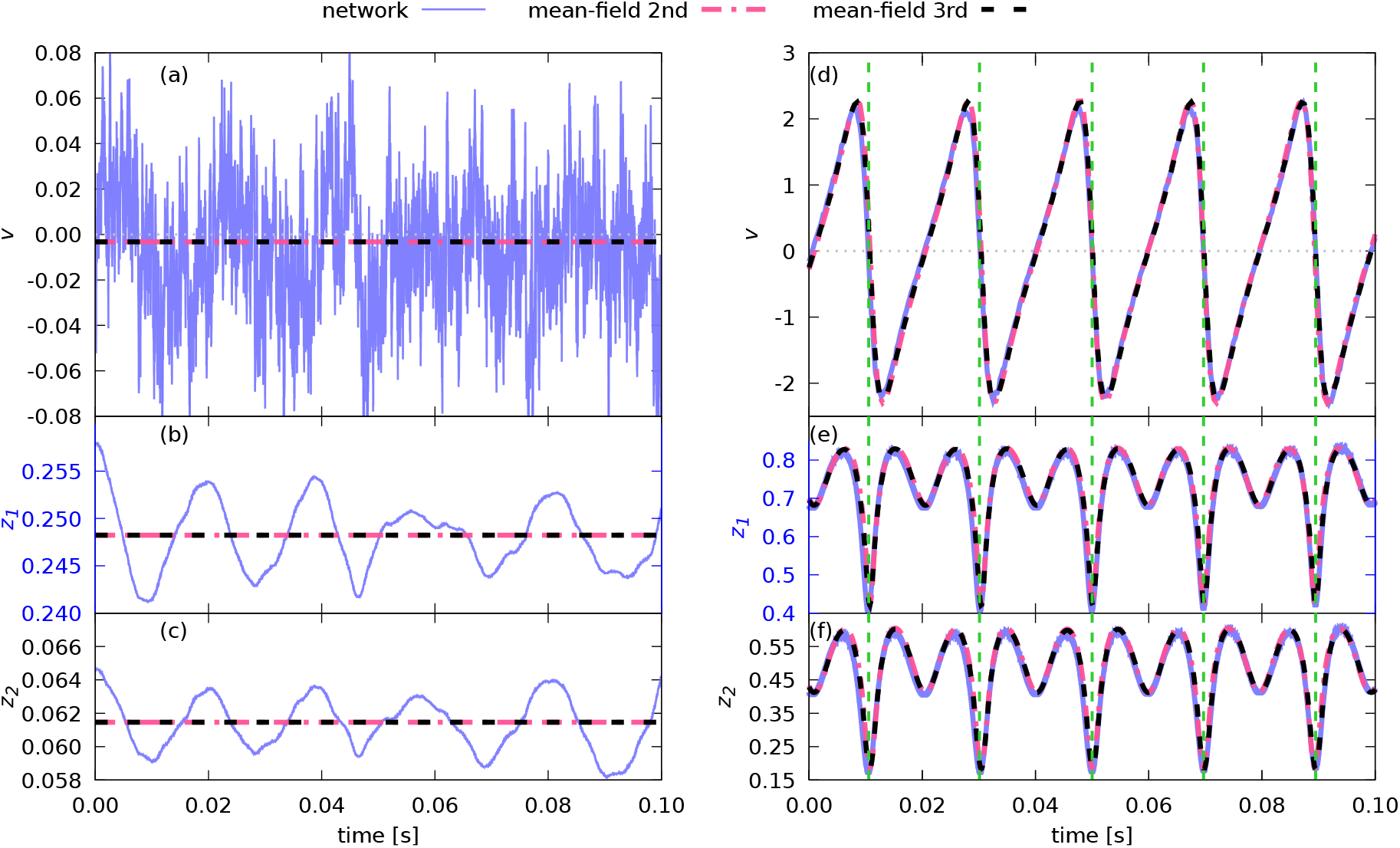
The mean membrane potential *v*(*t*) (a and d) and the Kuramoto-Daido order parameters *z*_1_ (b and e) and *z*_2_ (c and f) versus time. The data refer to the coexistence regime, namely *σ* = 0.00842, the asynchronous (clustered) state are reported on the left (right). Other parameters as in Fig. 1.

It is interesting to examine the level of synchronization in the two regimes as measured by the Kuramoto order paramters *z*_1_ and *z*_2_, see Fig. 4 (b,c and e,f). In the asynchronous state shown in panels (b,c) the neural mass results give a finite value for *z*_1_ and *z*_2_, while for an asynchronous regime one would expect zero values in the mean-field limit. The values of *z*_1_ and *z*_2_ obtained by the network simulations oscillate in an irregular fashion slightly around the mean-field value. For what concerns the clustered regime, the order parameters reveals periodic oscillations with the same period as *v*(*t*) and significant amplitudes. In this case the neural mass and the network results essentially coincide as shown in panels (e and f).

In order to understand the reason why *z*_1_ and *z*_2_ have a finite value in the asynchronous regime let us investigate the distribution of the phases as defined in Eq. (17). Histograms of these phases are shown in Fig. 5 for the asynchronous and clustered regime. In the asynchronous regime the phases are not equally distributed in [−*π*; +*π*] as expected they exhibit a peak around zero instead. This peak is much more pronounced in the clustered regime, but there is no evidence of the two clusters. This is due to the fact that the phase definition Eq. (17) is related to the membrane potential value, whose values also displays similar unimodal PDFs (see the Supplementary Material for animations of the phase histograms from Fig. 5 and the corresponding membrane potential histograms), and not to the firing time of the corresponding neuron, thus making this phase unsuitable to characterize the observed neural dynamics.

**Figure 5.**
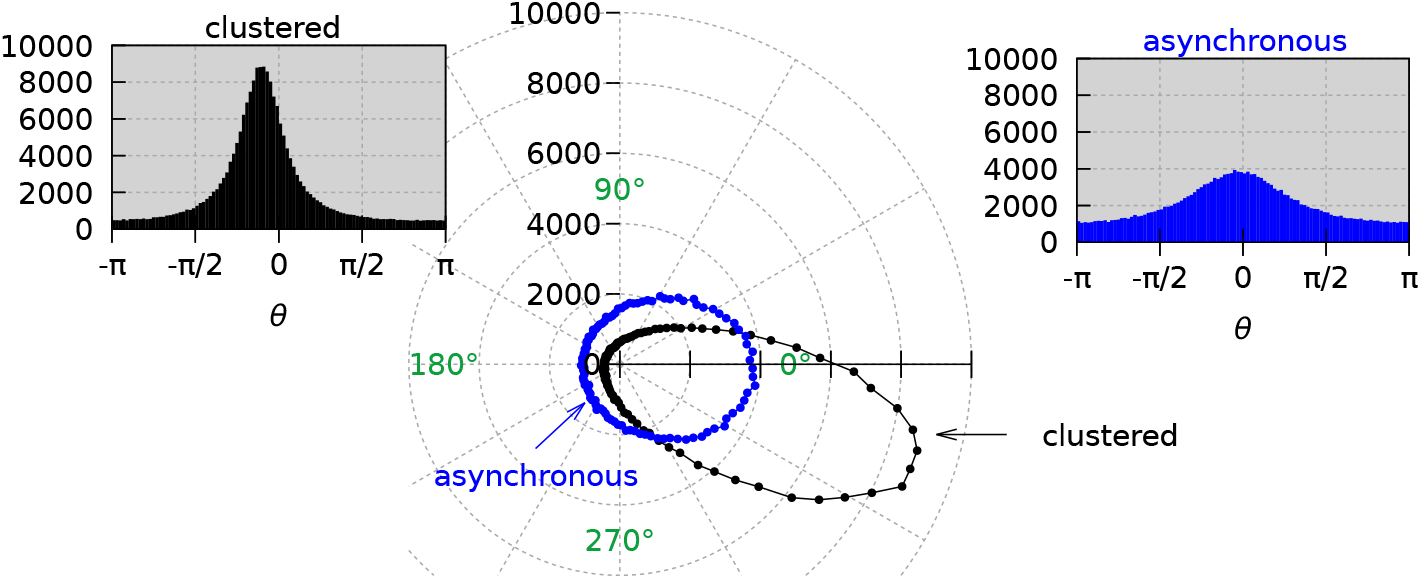
Snapshots of the phases as calculated via the expression (17) displayed as a polar plot for asynchronous (blue) and clustered (black) regimes. The insets show the histograms of the same data done over 100 bins. The data refer to a system size *N* = 200000 and *σ* = 0.00842. Other parameters as in Fig. 1.

Let us now consider the distributions of the phases as obtained by the firing times via the definition Eq. (19) for the asynchronous and oscillatory regimes. The results are shown in Fig. 6. Note that animations of these histograms can be found in the Supplementary Material. In the asynchronous case, as expected, the phases are uniformly distributed. The results for the oscillatory regimes reveal that the neurons are arranged in two clusters in phase opposition (at a distance *π*) from one another. In this case we expect that the Kuramoto order parameter 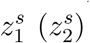 should be zero (order one) since the 2 clusters are in phase opposition.

**Figure 6.**
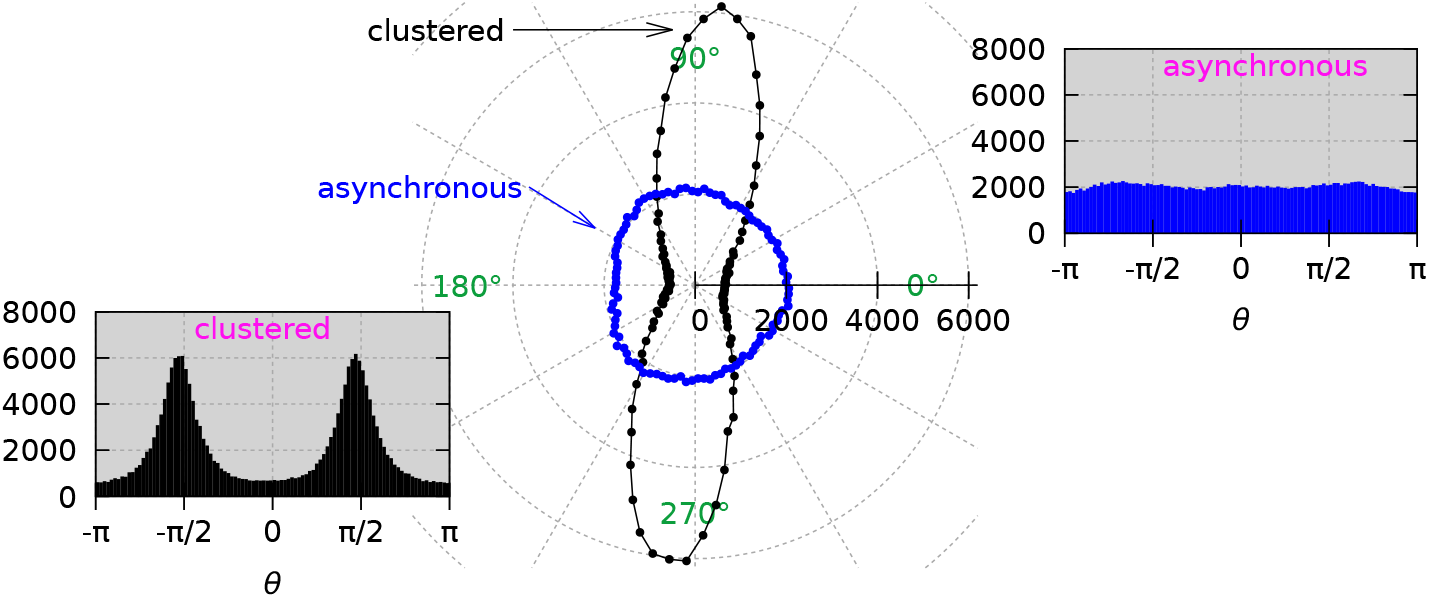
Snapshots of the phases as calculated via the expression (19) displayed as a polar plot for asynchronous (blue) and clustered (black) regimes. The insets show the histograms of the same data done over 100 bins. The data refer to a system size *N* = 200000 and *σ* = 0.00842. Other parameters as in Fig. 1.

To get some more insight on these two dynamical states, we will examine the raster plots as a measure of the microscopic network activity joined to the traces of the Kuramoto-Daido order parameters *z*_*k*_ and 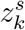 for the macroscopic counterpart. These are shown in Fig. 7.

**Figure 7.**
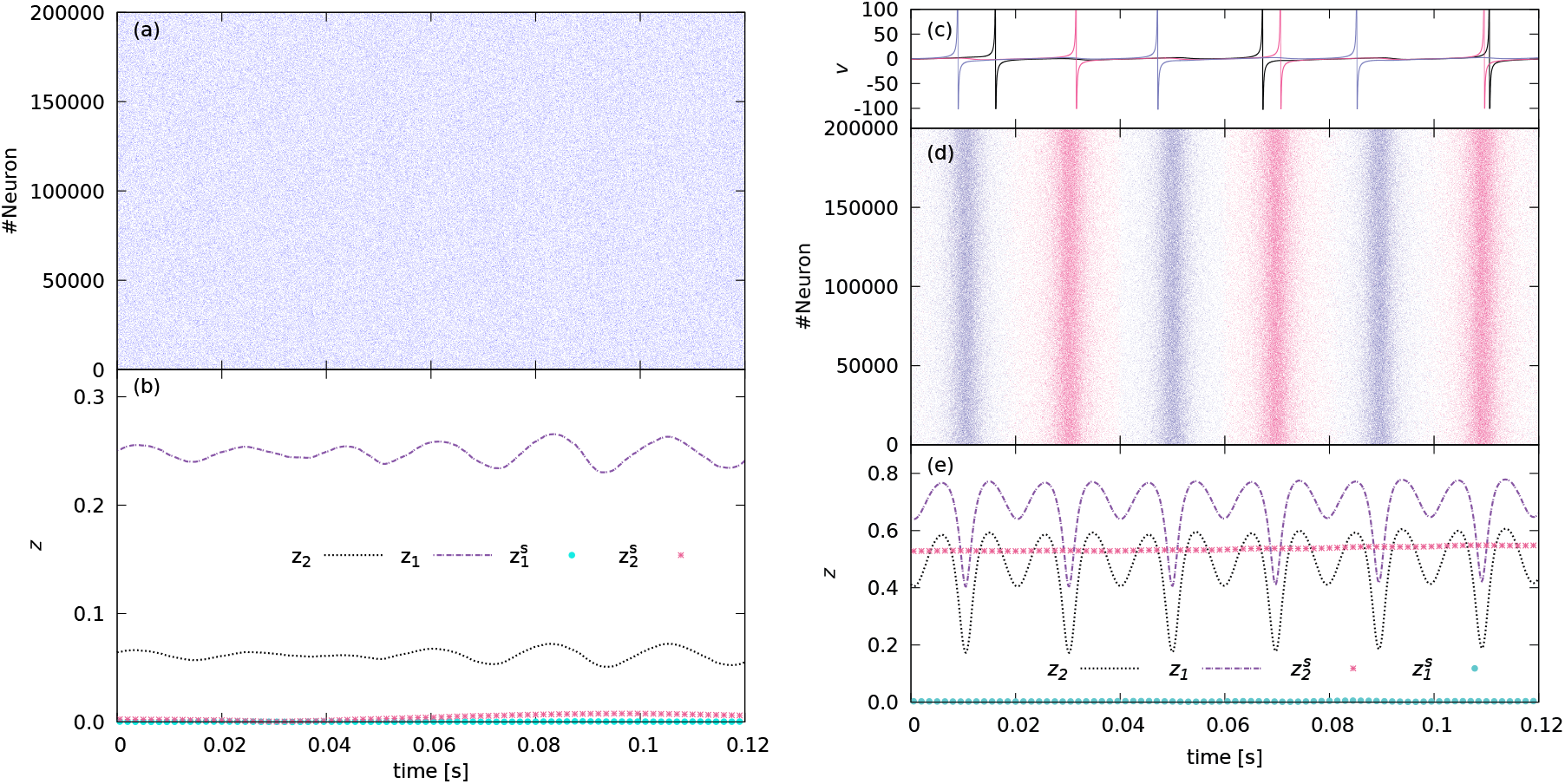
Raster plot (a and d) and the corresponding order parameters *z*_*k*_ and 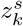 (b and e) versus time. In (c) the membrane potential of three generic neurons are displayed. The results for the asynchronous (clustered) regime are shown in the left (right) row. In the raster plot (d) the color of the dots indicates in which cluster the corresponding neuron is at time *t* = 0 based on their next spiking event. The data refer to a system size *N* = 200000 and *σ* = 0.00842. Other parameters as in Fig. 1.

In the asynchronous regime, the raster plot in panel (a) does not display any structure and the corresponding Kuramoto-Daido order parameters 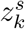, *k* = 1, 2 estimated by the firing-times are of 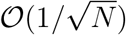 as expected (see panel (b)). As shown in panel (b) the values of *z*_1_ and *z*_2_ are instead definitely finite due to the fact that the phases obtained via the transformation Eq. (17) are not uniformly distributed, even in this regime.

In the clustered regime the raster plot (shown in panel (d)) reveals bursts of activity of the neurons interrupted by a low activity phase. In each population burst roughly 50% of the neurons participate. In the raster plot, the spiking times are visualized by red and blue colored dots based on which cluster the corresponding neuron belonged at time *t* = 0, for which we used the next spiking event of the corresponding neuron. In the time window reported in panel (d) the clusters are apparently stable, however on a longer run the two ensembles will mix up completely despite the macroscopic dynamics remaining always characterized by two equally populated clusters. To exemplify these behaviours in panel (c) the membrane potential traces for 3 characteristic neurons have been reported: the red (blue) neuron is always firing within the red (blue) burst, while the black one is initially firing within the blue burst but then it skips 2 population bursts and finally joins the red burst.

In panel (e) we report the corresponding Kuramoto-Daido order parameters versus time, the parameters *z*_1_ and *z*_2_ display a periodic behaviour and attain the minimum value whenever a burst occurs, due to the repulsive nature of the couplings. On the other hand, 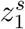 stays always close to zero as expected for two phase clusters in phase opposition, while *z*_2_ has a constant finite value larger than 0.5 indicating that the composition of the clusters is stable in time.

#### 4.1.3. Stability of the asynchronous regime

In this Paragraph we will analyze the stability of the asynchronous state for increasing noise amplitude. For this we will employ the stability indicator *ρ* = *ρ*(*γ*) introduced in Eq. (24) as a function of the parameter *γ* controlling the initial distribution of the membrane potentials according to Eq. (23). The value *γ* = 1 (*γ* = 0) corresponds to an initialization of the neurons with membrane potentials distributed according to the LD expected for the asynchronous case (with identical values of the membrane potentials *V*_0_).

We have estimated the stability indicator *ρ*(*λ*) for a system size *N* = 32000 (due to cpu limits) and the results are reported in Fig. 8 for different noise amplitudes. For the noise amplitude *σ* = 0.0025, which is smaller than *σ*_*SN*_ ≃ 0.004 and therefore outside the coexisting region, we observe that the asynchronous state is stable for any *γ*-values, as expected. For larger noise amplitude *σ > σ*_*SN*_, large perturbation of the asynchronous distribution, as measured by 1 − *γ*, can induce transitions towards the clustered regime with some finite probability.

**Figure 8.**
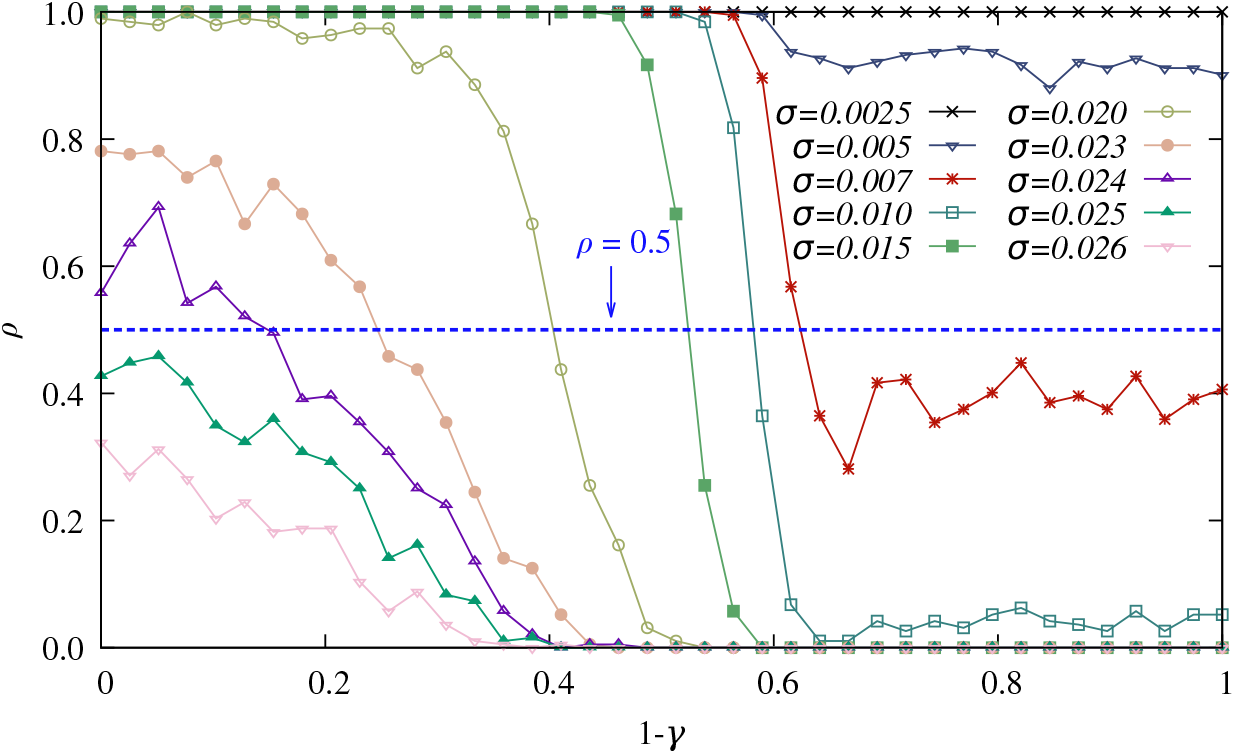
Stability indicator *ρ* versus 1 − *γ*, where *γ* is the factor entering in (23). For each measurement of *ρ* we considered *M* = 192 different realizations and each time we estimated Σ_*v*_ from a time series of *T*_*W*_ = 0.3 s after discarding a transient *T*_*t*_ = 16 s to evaluate if the system remains asynchronous or becomes clustered. The dashed line indicates *ρ* = 0.5, i.e., where the system has an equal probability to move towards either the asynchronous or the clustered state. The data refer to a system size *N* = 32000, other parameters as in Fig. 1.

At noise amplitudes *σ* ≥ 0.015, for sufficiently synchronized initial conditions, namely *γ <* 0.4, the system has a probability of almost 100% to leave the asynchronous state, thus indicating a clear coexistence of the 2 regimes. For *σ* ≥ 0.023 (i.e. in proximity of the Hopf bifurcation identified in the mean-field formulation *σ*_*H*_ = 0.0243) the probability to stay in the asynchronous case is smaller than 100% even for the unperturbed initial conditions, corresponding to *γ* = 1, in this case we expect that by simulating for a longer time period *T*_*t*_ we would actually measure *ρ* = 0.

In order to identify the critical noise amplitude above which the asynchronous state is unstable, we measure the value 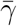 for which 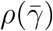 crosses 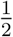 for various noise amplitudes *σ*. Thus, this indicates that for 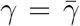 one has 50% of probability to end in the asynchronous or in the clustered state. For this we estimated the indicator with more precision in proximity of 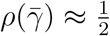 To this end we considered *M* = 384 realizations of the initial perturbed state for each value of *γ*, and we estimated via an interpolation the value 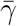. In particular, we expect that the standard deviation of *ρ*(*γ*) will be maximal at exactly one half, therefore we fitted such standard deviation to a Gaussian curve for different *γ* value and we extrapolated the value 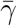 where the curve attains its maximum.

In Fig. 9 we show the obtained values 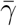 as a function of noise amplitude *σ* (violet crosses), we observe that 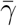 grows with the noise amplitude and approaches the value 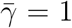. To identify the critical noise amplitude *σ*_*c*_ above which the system always ends up in the clustered regime for any *γ* value we have fitted the numerical data with this function:

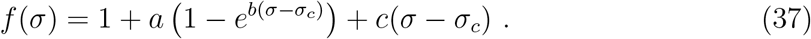

**Figure 9.**
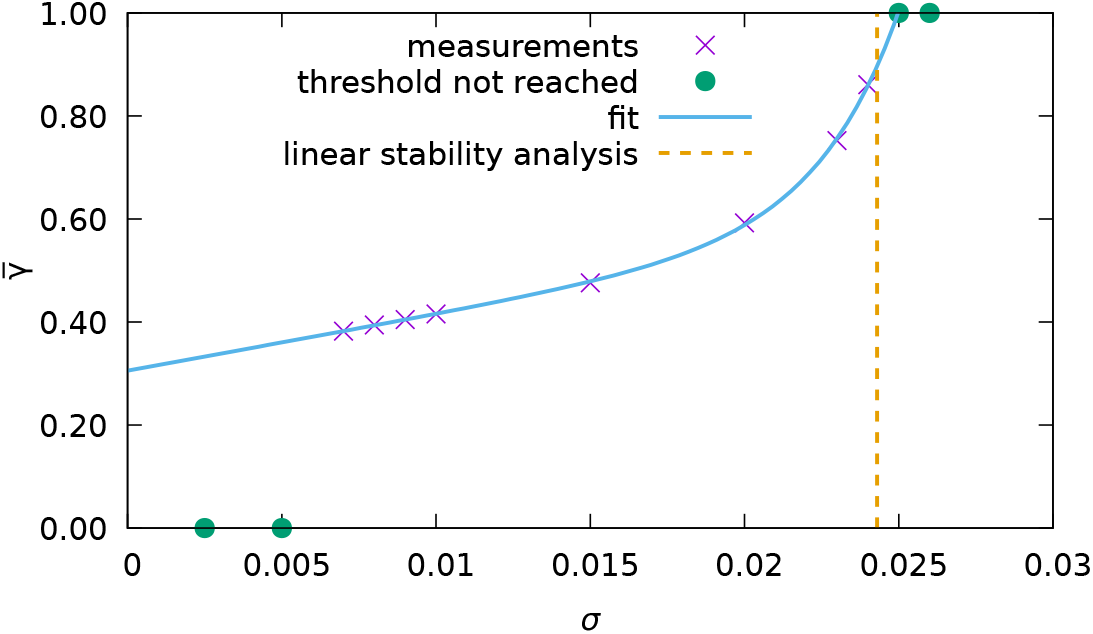
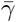 versus the noise amplitude *σ*. The green circles represent *σ* values for which *ρ*(*σ*) = 0.5 was never reached. The blue solid line is a fit to the data (violet crosses) performed via the expression (37), the fit is employed to extrapolate the critical noise value *σ*_*c*_ for which 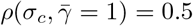. The vertical dashed line shows the mean-field prediction *σ*_*H*_ for the onset of COs. Errorbars are omitted since the errors are smaller than symbol size. Other parameters as in Fig. 8 apart the number of different realizations, that now is *M* = 384.

By excluding from the fit the data where the threshold value *ρ* = 0.5 was not reached (green points) we obtain the following parameter values *a* = −0.42(1), *b* = 376(28), *c* = 10.9(8) and *σ*_*c*_ = 0.02498(6). As you can see the fit works pretty well. The extrapolated critical value of the noise is in quite good agreement with the mean-field result obtained from the linear stability analysis of the asynchronous state that indeed was *σ*_*H*_ = 0.0243, the difference on the third significative digit can be due to finite-size and nonlinear effects.

In summary the new method here introduced to study the stability of the asynchronous regime works reasonably well when compared with the linear stability analysis, that in the present case is feasible due to the existence of low-dimensional mean-field formulations, but usually in a high dimensional network is quite difficult to implement. Therefore, this new method can represent an useful alternative to the linear stability analysis and it can find applications in many complex network systems. Furthermore, it gives also information concerning the basins of attraction of the two regimes in the coexistence region, that the linear stability analysis is unable to provide.

#### 4.1.4. Characterization of the clustered dynamics

In this Paragraph we would like to examine the clustered dynamics of the neurons in more details. In particular, we wish to characterize the erratic behaviours that lead the neurons to deviate from a perfectly locked evolution, where the neurons fire every second burst.

We will first examine the evolution in time of the fraction of surviving neurons *S*(*t*) (or survival probability). As shown in Fig. 10, *S*(*t*) has an initial decay on the interval [0 : 30] s very well described by the following function

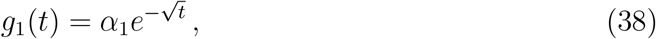

with *α*_1_ = 1.65(1). The initial decay of *S*(*t*) is followed at later times by an exponential tail of the form

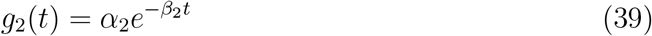

with *α*_2_ = 0.142(2) and *β*_2_ = 0.1026(3) Hz. The functions *g*_1_ and *g*_2_ are part of the same class of survival probabilities associated to the so-called Weibull PDF [45]

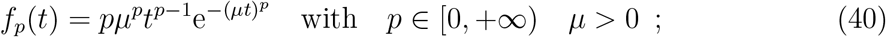

where *g*_1_ (*g*_2_) corresponds to *p* = 1*/*2 (*p* = 1). For *p <* 1 the failure rate, the rate to emit a spike in an irregular manner, decreases over time, since the neurons that displays an irregular spiking are eliminated from the population of the regular spiking ones. The neurons remaining after this initial phase have a failure rate *β*_2_ that is constant over time, since their survival probability has an exponential profile, which typically emerges due to some underlying random Poissonian process.

**Figure 10.**
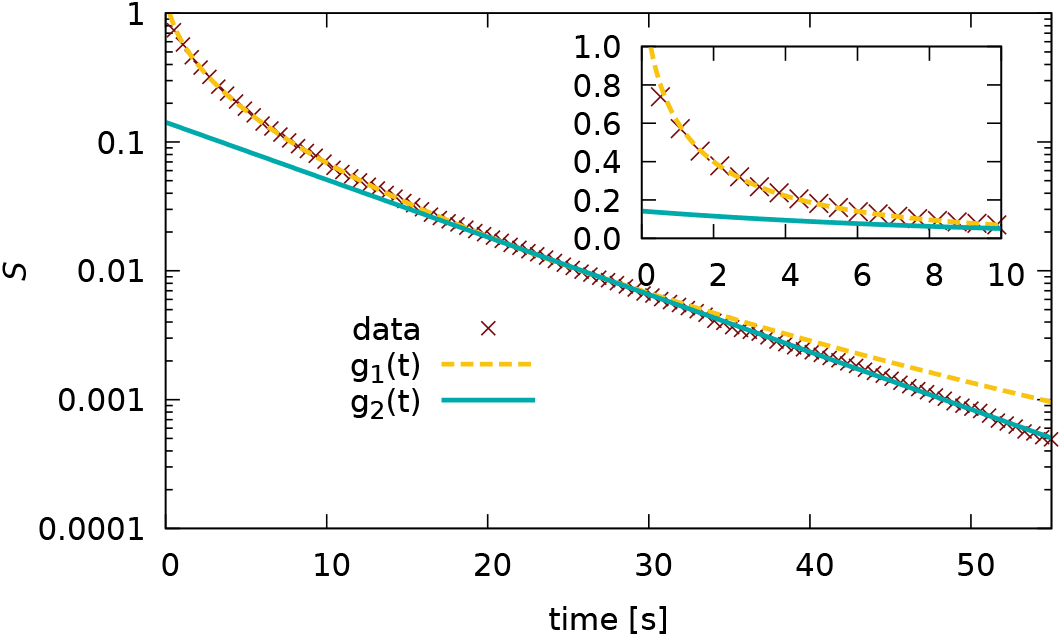
Fraction of surviving neurons *S*(*t*) versus time in a semi-logarithmic scale. We fitted the data to the function *g*_1_(*t*) (38) in the time interval [0 : 30] s and to the function *g*_2_(*t*) (39) in the interval [20 : 55] s. The inset shows the evolution over the first 10 seconds in a linear scale. In this case we have identified *N*_none_ = 467 silent neurons. The data refer to a system size *N* = 200000 and *σ* = 0.00842 for the clustered regime. Other parameters as in Fig. 1.

As we will see in the following for a homogeneous network *S*(*t*) is very well described by Eq. (39) over the whole time interval. Thus suggesting that this decay is likely due to the action of the noise injected in the system, since this is the only source of irregularity in the homogeneous case. While the initial decay described by the function *g*_1_(*t*) should be related to the heterogeneous distribution of the synaptic couplings. In summary, initially the neurons displaying failures in their periodic activity are the ones with *J*_*i*_ sufficiently different from *J*_0_, while the successive decay involves neurons with coupling in proximity of *J*_*i*_ = *J*_0_. This aspect will be further analyzed in the following, where we will correlate in more details the irregular evolution of the *i*-th neuron to its synaptic coupling *J*_*i*_.

Next, we examine the PDF *P* (*λ*) of the fraction *λ* of irregular spikes (28) emitted by each neuron. We report *P* (*λ*) in Fig. 11 for increasing duration of the simulations and therefore for an increasing number of population bursts.

**Figure 11.**
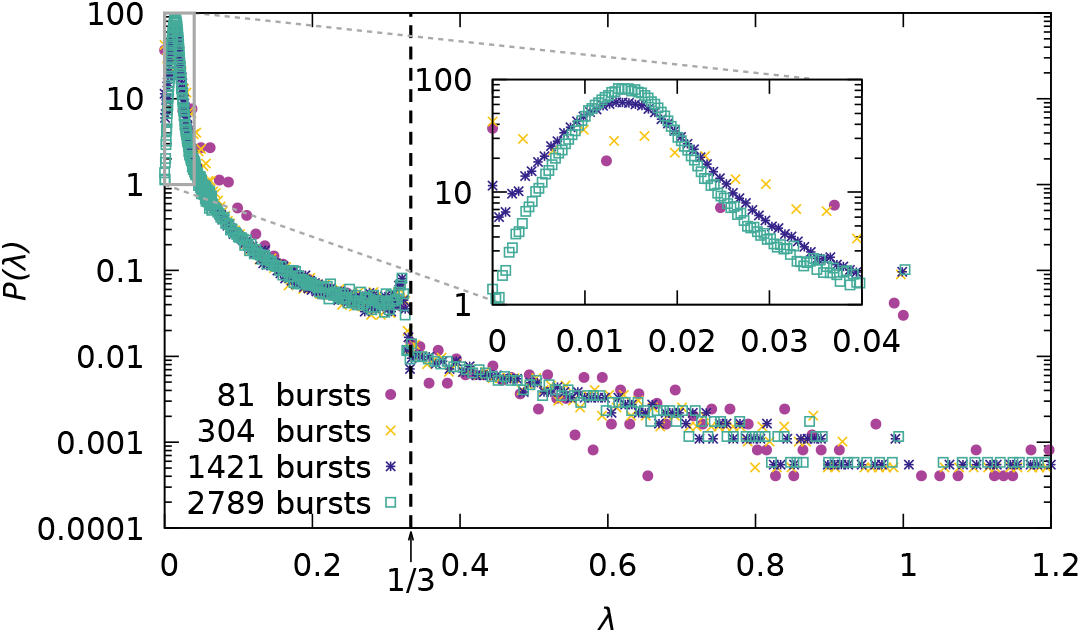
Probability distribution function *P* (*λ*) of the fraction of irregular spikes *λ* for different time durations in semi-logarithmic scale. The inset shows a zoom around the main peak. In this case we have removed from the estimation of the PDF *N*_none_ = 467 silent neurons. The data refer to a system size *N* = 200000 and *σ* = 0.00842 for the clustered regime. Other parameters as in Fig. 1.

From the figure it is evident that the PDF is converging to a limiting profile for longer duration of the measurements. This asymptotic shape reveals a peak around *λ* ≃ 0.0145 corresponding to the neurons with couplings in proximity of *J*_*i*_ = *J*_0_ = −20, i.e. to the maximum of the LD of the synaptic coyplings. Furthermore, *P* (*λ*) reveals a clear discontinuity at *λ* = 1*/*3, whose origin will become clear in the following.

Let us now characterize in details how the fraction of irregularly emitted spikes of neuron *i* depends on its synaptic coupling *J*_*i*_. To this aim we have estimated *λ*_*i*_, 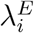 and 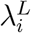 for each neuron as well as the fraction *ν*_*i*_ of emitted spikes with respect to the total number of population bursts (*ν*_*i*_ in absence of irregularity should be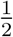).

These quantities are shown versus the corresponding *J*_*i*_ as scatter plots in Fig. 12. It is important to notice that the scatter plots actually look like smooth functions for all the considered indicators, not much “scatter” visible. This suggests the existence of a functional relationship between the measure values and the value *J* of the synaptic coupling.

**Figure 12.**
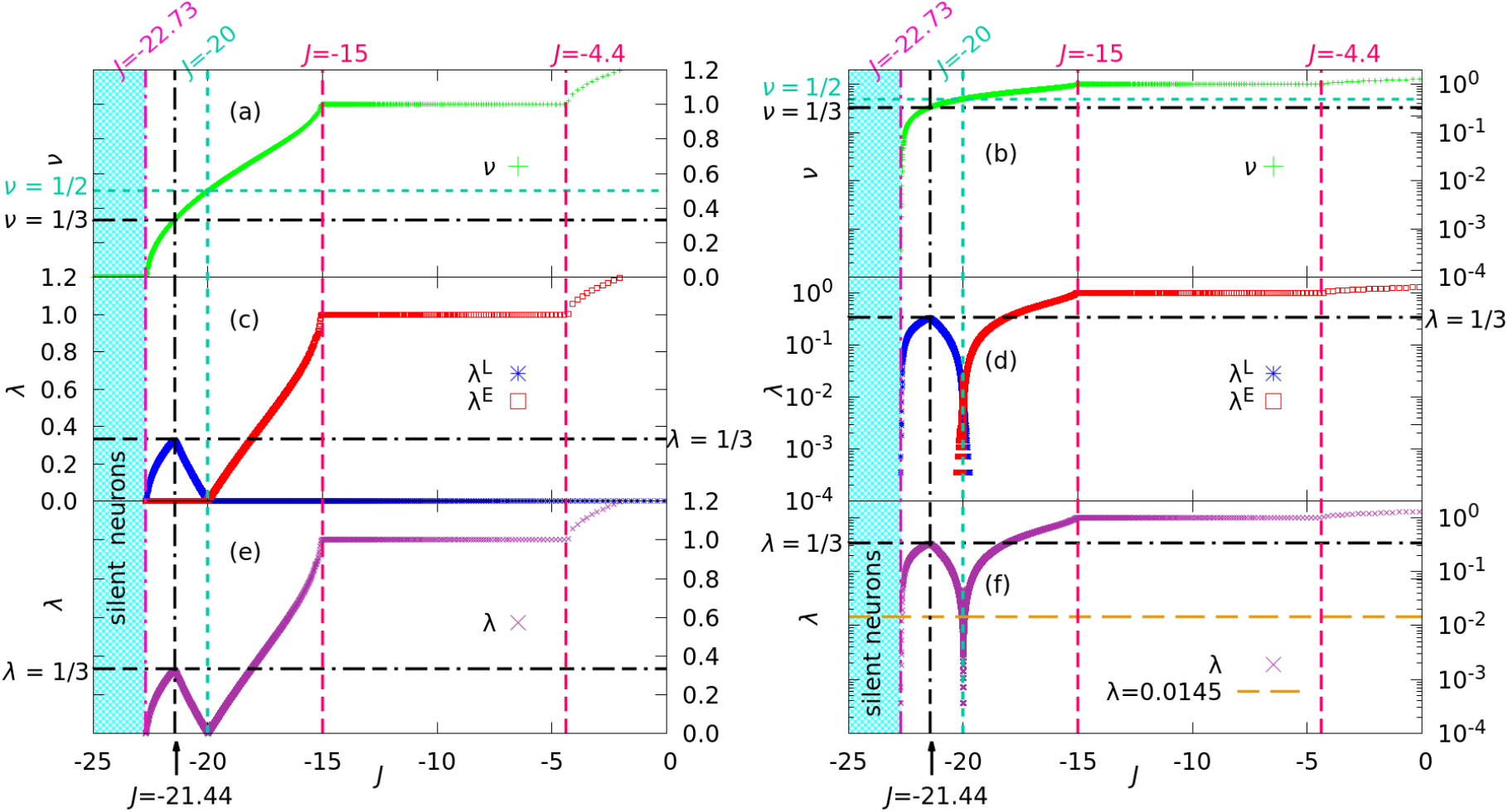
Scatter plots for *ν*_*i*_ (a and b) and for the indicators *λ*_*i*_ (e and f), 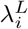 and 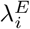 (c and d), measuring the fraction of irregular spikes, versus the respective synaptic coupling *J*_*i*_. The left panels are in linear scale, while the ones on the right side are the same data reported in semi-logarithmic scale. The measurements corrrespond to the ones reported in Fig. 11 for a total of 2789 bursts.

For sufficiently small *J*_*i*_ ≤ −22.73 the neurons appear to be silent on the considered integration time scale. As shown in panels (a and d), *ν*_*i*_ is growing with the synaptic coupling, apart in the locking regions. On the contrary, 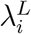 and *λ*_*i*_ have a non monotonic dependence on *J*_*i*_, with a maximum at *J*_*i*_ = −21.44 where these parameters reach the value of exactly 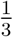. *λ*_*i*_ displays a minimum at *J*_*i*_ = *J*_0_ = −20, where it attains extremely small values (see panels (c and d) and (e and f)). For larger *J*_*i*_ essentially 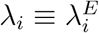 and they are both increasing with *J*_*i*_, again apart the locking intervals. As a general remark the irregularity in the periodic firing of the neurons is due to early (late) delivered spike for *J*_*i*_ *< J*_0_ (*J*_*i*_ *> J*_0_).

Let us now try to understand the non monotonic behaviour of *λ*. The local maximum 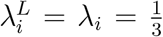 at *J*_*i*_ ≤ −22.73 corresponds to 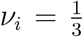, which means that the corresponding neurons fire very regularly at every 3rd burst. The origin of the maximum is due to the fact that for smaller *J*_*i*_ the neurons are firing less and less, thus the value of 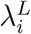 and *λ*_*i*_ should necessarily decrease, however for larger *J*_*i*_ the two parameters are also decreasing. This is due to the fact that the regular behaviour occurs whenever the neurons fire exactly every two spikes, and this state is approached by increasing *J*_*i*_ towards *J*_0_.

Indeed, for *J*_*i*_ = *J*_0_ the rate is exactly 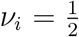 and at the same time 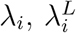 and 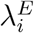become quite small and close to zero (see the semi-logarithmic plots in panel (d)). In particular, from panel (f) it is evident that the neurons contributing to the peak of *P* (*λ*) reported in Fig. 11 are those with *J*_*i*_ = *J*_0_, see the dashed orange line indicating the value *λ* = 0.0145, where *P* (*λ*) attains its maximum.

It is interesting to notice that in the interval *J* ∈ [−15, −4.4] we have a perfect locking of the activity of these neurons with the population bursting since *ν* = 1, and unsurprisingly, also *λ*^*E*^ = 1. The locking region resembles an Arnold tongue 1:1, for *J >* −4.4 the locking is lost and *ν* and *λ* continues to increase. We have indications that another locking region with *ν* = 2 emerges at quite large *J*, around 25 ≤ *J* ≤ 29, however our system size is too small to have a good statistics there.

Let us now come to the explanation of the discontinuity observed in Fig. 11 for the PDf *P* (*λ*) at 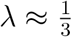. This is due to the fact that the maximal value of *λ*_*L*_ is 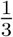, thus for 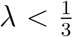 to the *P* (*λ*) contribute both early and late delivered spikes, while for 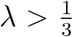 the contribution to the irregularity is due only to early delivered spikes.

To get further insight on the microscopic dynamics induced by the synaptic couplings, we define a global phase Φ similar to Eq. (19). However, instead of the spike times of the individual neurons we consider here the population burst times *b*_*k*_:

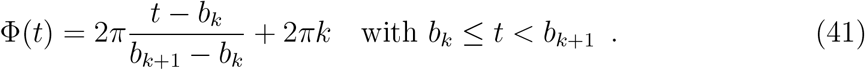

Moreover, Φ(*t*) can be employed to characterize the activity of the *i*-th neuron with respect to the network activity by defining the global phase difference associated to two successive spikes of neuron *i*:

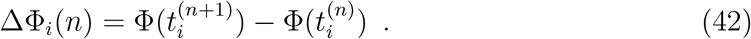

In Fig. 13 we report the average and variance of ΔΦ_*i*_ estimated over all the spike times of neuron *i* versus the corresponding synaptic coupling *J*_*i*_. A clear functional relationship emerges also in this case. As expected, the value of the mean of ΔΦ_*i*_ is 4*π* for *J*_*i*_ = −20 indicating that the neurons fire every second burst. For *J*_*i*_ *< J*_0_ (*J*_*i*_ *> J*_0_) ΔΦ_*i*_ grows (decreases) indicating that the neurons fire slower (faster). Moreover, the phase locking at ΔΦ_*i*_ = 2*π* for neurons with *J*_*i*_ ∈ [−15, −4.4] is also evident from panel (a).

**Figure 13.**
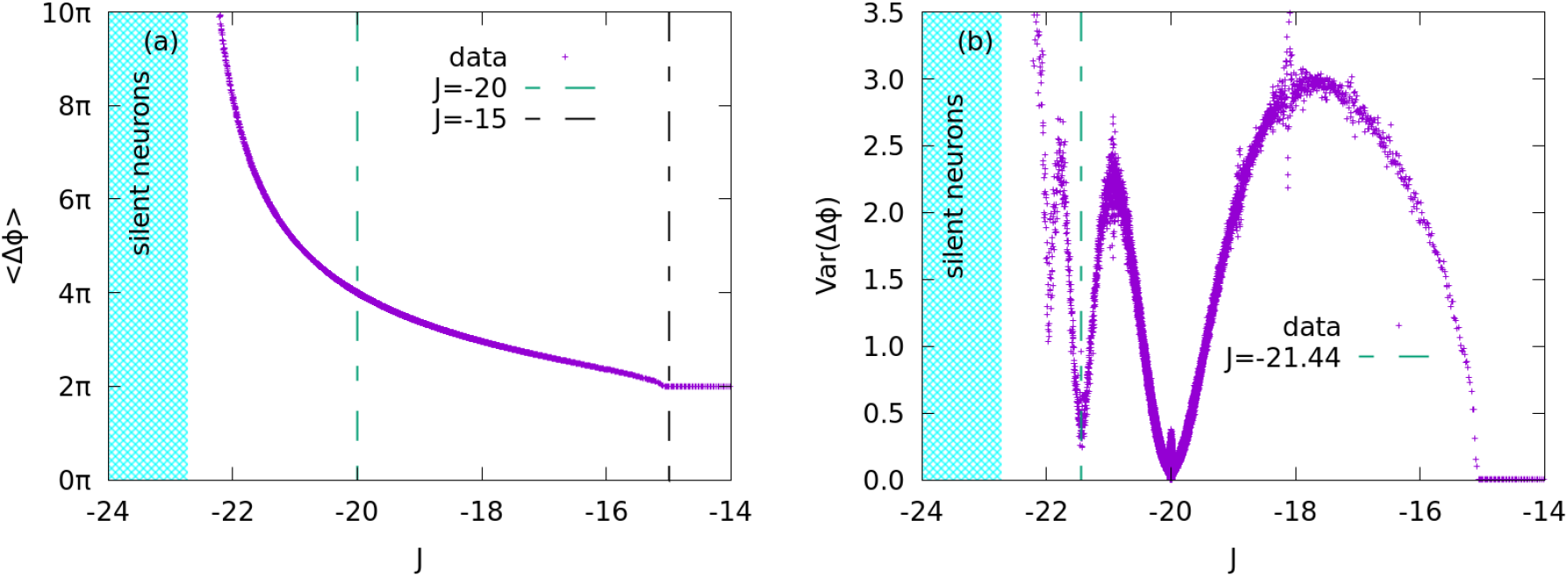
Mean (a) and variance (b) of the global phase difference ΔΦ_*i*_ for each neuron reported as a scatter plot versus the correpsonding coupling *J*_*i*_. The data have been obtained from the same measurements employed in Fig. 11.

The analysis of the variance of ΔΦ_*i*_ reveals more interesting aspects. A variance close to zero suggests that the corresponding neuron fires very regularly, i.e., basically with a constant firing rate. In contrast a high variance indicates a distribution of the global phase differences ΔΦ_*i*_ exhibiting more peaks. As shown in Fig. 13 (b), the variance attains its minimal value for *J*_*i*_ = −*J*_0_ for the regular firing neurons, for which ⟨ΔΦ_*i*_⟩ ≃ 4*π*. Moreover, minima in the variance are observable also whenever ⟨ΔΦ_*i*_⟩ ≃ 6*π* at *J* ≈ −21.44, and also at lower *J*_*i*_ where ⟨ΔΦ_*i*_⟩ ≃ 8*π*. Furthermore, the variance vanishes in the locking region (*J* ∈ [−15, −4.4]) where ⟨ΔΦ_*i*_⟩ = 2*π*. The maxima in the variance are instead observable when ⟨ΔΦ_*i*_⟩ ≃ (2*k* + 1)*π* for *k* = 0, 1, 2. For the case ⟨ΔΦ_*i*_⟩ ≃ 3*π*, we observe that the corresponding neurons emit two spikes almost in correspondance with two successsive population bursts and then skip one burst. This amounts to a sequence of phase differences ΔΦ_*i*_ = 2*π*, 4*π*, 2*π*, 4*π*, …, that gives an average global phase of 3*π* and a distributions of the phases with two equally relevant peaks and thus to a high variance.

We can safely affirm that the neurons tend to fire in correspondance of the bursting activity of the network, in general every two bursts, but as shown above they can present more complex combinations of locking *n* : *m* with the population bursting.

### 4.2. Homogeneous synaptic couplings

As we have seen in Subsection 3 for the case with homogeneous couplings, i.e., Δ_*J*_ = Δ_*η*_ = 0, the linear stability of the mean-field model predicts that the asynchronous state is unstable whenever the noise amplitude is finite. However, the mean-field approach is no more strictly valid in the fully homogenous case, since the Ott-Antonsen manifold is no more attractive in such a case [46]. Therefore, we will limit to network simulations in order to numerically investigate the stability of the asynchronous state as well as possible coexistence regime with a collective oscillatory dynamics.

#### 4.2.1. The clustering transition

To this aim we performed quasi-adiabatic simulations of the QIF network by varying the noise amplitude *σ* and by evaluating the variance Σ_*v*_ of the mean membrane potential *v*, analogously to the analysis done in the heterogeneous case, whose results have been reported in Fig. 3. In the present case, by increasing adiabatically *σ* in the interval [0, 0.015] we observe that the asynchronous regime appears to remain stable up to noise amplitude *σ*_*c*_ ≃ 0.011, while for larger noise COs emerge. Successively, by decreasing *σ* the oscillatory regime remains stable down to *σ* = 0, thus we have a coexistence regime in the whole interval *σ* ∈ [0, 0.011], as shown in Fig. 14. Note the contrast with the heterogeneous case where, in the absence of noise, only the asynchronous regime was observable.

**Figure 14.**
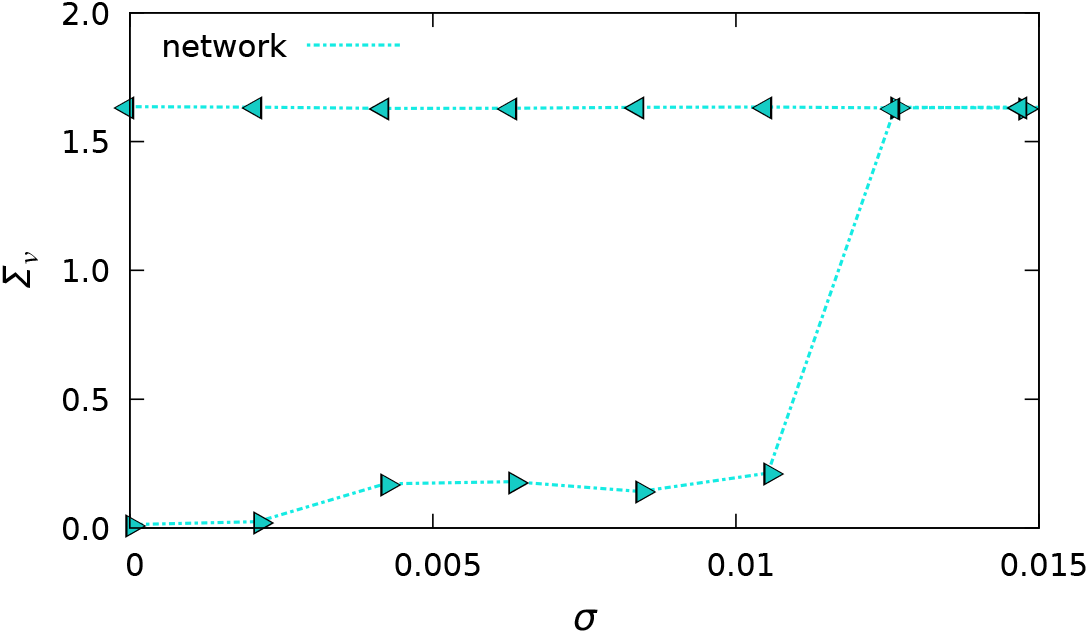
Variance Σ_*v*_ of the mean membrane potential *v* versus noise amplitude *σ* obtained via quasi-adiabatic simulations. The decrease or increase of the noise amplitude *σ* performed during the adiabatic simulations is indicated by the direction of the triangles’ tip. The dashed lines are simply intended as a visual aid. The parameters for the quasi-adiabatic simulations are 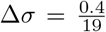, *t*_*T*_ = 20 s, *t*_*S*_ = 25 s. The other parameters are set as in Fig. 3 with Δ_*J*_ = 0 for a network size *N* = 200000.

The oscillatory regime is once more a clustered regime, where the neurons fire in population bursts and each burst involves almost half of the population.

#### 4.2.2. Stability of the asynchronous regime

Let us now analyze, how the stability of the asynchronous state depends on the noise amplitude. To perform this analysis we have employed (as in Subsection 4.1.3) the indicator *ρ* = *ρ*(*γ*) introduced in Eq. (24) as a function of the parameter *γ*. The corresponding results are reported in Fig. 15 for various noise amplitudes *σ* ∈ [0, 0.015].

**Figure 15.**
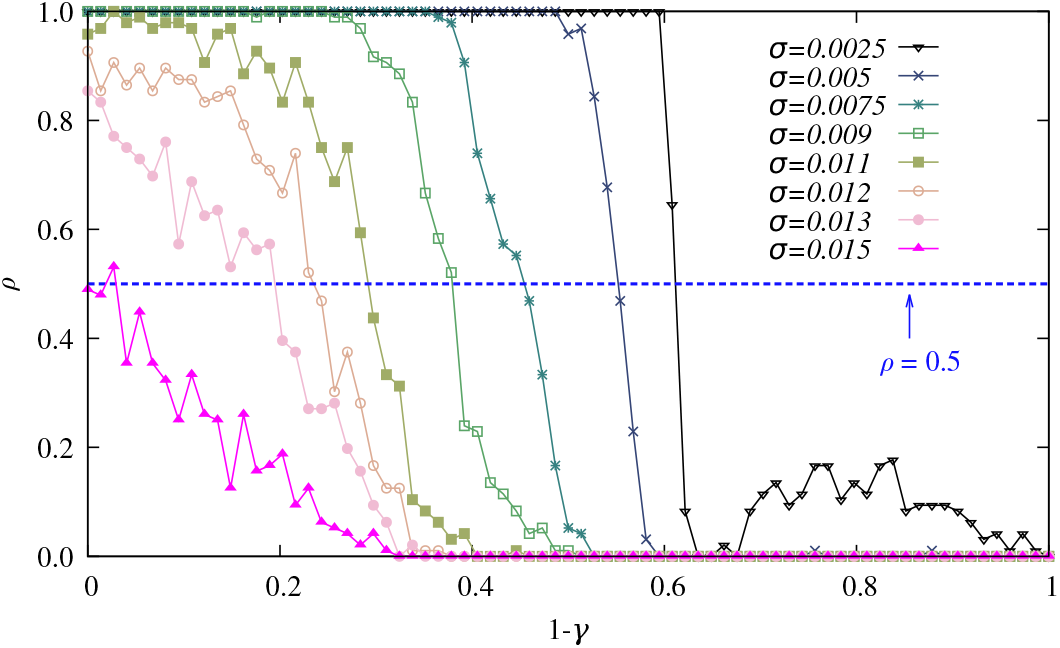
Stability parameter *ρ* versus 1 − *γ*, where *γ* is the modulation factor entering in (23). For each measurement of *ρ* we averaged over *B* = 96 different noise realizations and each time we estimated Σ_*v*_ from a time series of duration *T*_*W*_ = 0.3 s after discarding a transient *T*_*t*_ = 16 s. The dashed line indicates *ρ* = 0.5, i.e., where the system has an equal probability to end in the asynchronous or clustered state. The data refer to a system size *N* = 32000, other parameters as in Fig. 14.

The main difference with respect to the heterogenous case, is that now even for the smallest noise amplitude considered, namely *σ* = 0.0025, for a sufficiently large distortion of the LD (namely, *γ >* 0.4) will destabilize the asynchronous state. This is a further confirmation that the clustered state is always stable, as already shown in Fig. 14. For increasing noise amplitudes the asynchronous state gets destabilized for larger and larger *γ* values and for *σ >* 0.011 even for *γ* = 1. Thus indicating that this is the critical noise amplitude above which only the clustered regime can be observed on the long-time limit.

In analogy to what done in Subsection 4.1.3, to better characterize this transition, we will estimate for various noise amplitudes the value 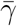 for which 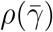 crosses 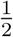 . In particular, we measured 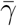 by using *B* = 192 to obtain a better accuracy. The results are displayed in Fig. 16, by performing a fit of the data to the function (37) we obtained the following parameter value *a* = −0.21(6), *b* = 376(205), *c* = 33(6) and *σ*_*c*_ = 0.0149(3). Thus obtaining a critical noise amplitude consistent with the previous estimations.

**Figure 16.**
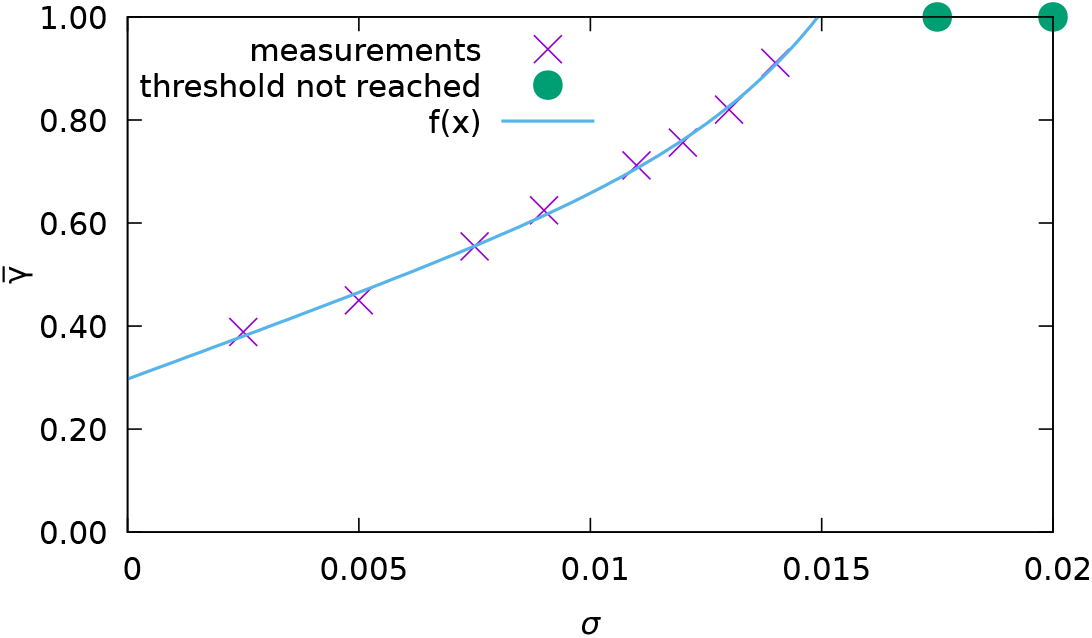
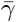 versus the noise amplitude *σ*. The green circles represent *σ* values for which *ρ*(*σ*) = 0.5 was never reached. The blue solid line is a fit to the data (violet crosses) performed via the expression (37), the fit is employed to extrapolate the critical noise value *σ*_*c*_ for which 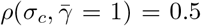 Other parameters as in Fig. 15 apart *B* = 192.

#### 4.2.3. Characterization of the clustered dynamics

Analogously to the heterogenous case the oscillatory dynamics is characterized by neurons firing alternately in the two population bursts. To measure the irregularity in this dynamics, we have examined, as in the heterogenous case, the fraction of surviving neurons *S*(*t*) defined as in Eq. (29), for the same noise amplitude *σ* = 0.00842. We now observe that *S*(*t*) can be well reproduced by a simple exponential decay Eq. (39) with parameters *α*_2_ = 0.853(2) and *β*_2_ = 0.0752(6) Hz, see Fig. 17. The mean lifetime of the periodic regular regime is now of the order of 13.3 s, definitely longer than in the heterogenous case. Since the only source of irregularity is now the noise, we confirm that the emergence of the irregularities in the firing process follows a Poissonian process. Furthermore, in contrast with the heterogeneous case we we did not observe any silent neurons, since none of the neurons receive very large inhibitory post-synaptic potentials, because the amplitude of the synaptic weight is the same for all neurons.

**Figure 17.**
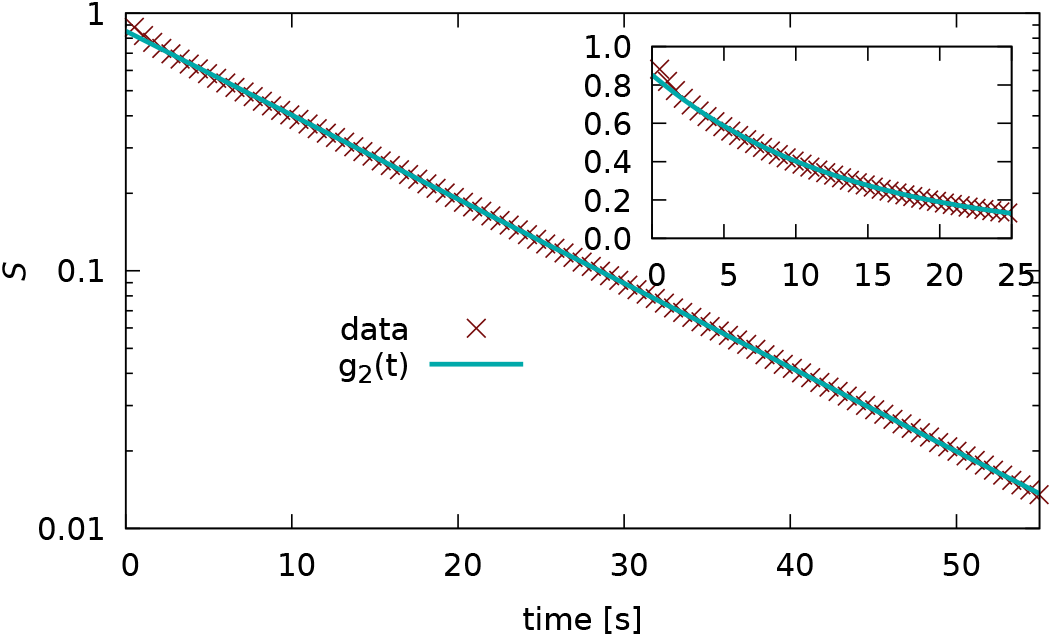
Fraction of surviving neurons *S*(*t*) verus time in semi-logarithmic scale. We also include a fit to the numerical data with the function *g*_2_(*t*) (39) (blue line). The inset shows the first 25 seconds of the evolution of *S*(*t*) in a linear scale. The parameters are fixed as in Fig. 14 for a noise amplitude *σ* = 0.00842 and we refer to the clustered phase.

Next, we have estimated *P* (*λ*), i.e. the PDF of the fraction *λ* of irregular spikes obtained for each neuron, this is reported in Fig. 18. We have integrated the network for the the same time duration as in Fig. 11, however due to the slightly higher frequency of the COs measured in the homogeneous case we observe more bursts.

**Figure 18.**
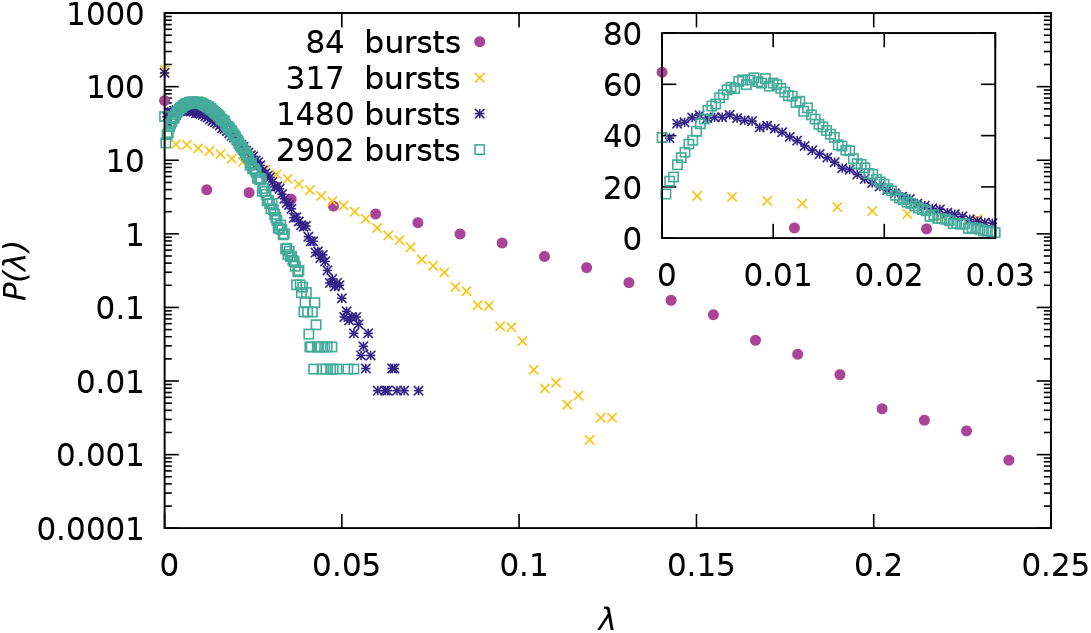
PDF *P* (*λ*) of the fraction of irregular spikes *λ* for different measurement durations in semi-logarithmic scale. The inset shows a zoom in linear scale. The parameters are fixed as in Fig. 14 for a noise amplitude *σ* = 0.00842 and we refer to the clustered phase.

In this case the PDF for *λ*^*E*^ and *λ*^*L*^ are identical (not shown) and this is clearly due to the absence of heterogeneity in the network. The noise induces with equal probability irregularities due to early or late spiking.

For increasing duration of the measurements we observe that *P* (*λ*) tends to get more and more localized around the maximum located at *λ* = *λ*_0_ ≈ 0.01. Due to the central limit theorem we expect that the function *P* (*λ*) has a Gaussian profile (limited to *λ >* 0) with a standard deviation *σ* scaling as 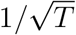, where *T* is the time duration. Indeed, by considering only values of *λ > λ*_0_ we have verified that ln 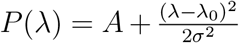 with *σ* ∝ *T*^*ξ*^ where *ξ* = −0.52 ± 0.02 for the data reported in Fig. 18.

### 4.3. Emergence of γ-oscillations in the clustered state

The clustered state is characterized by population bursts, corresponding to COs. It is of extreme interest to understand in which frequency range these oscillations occur. In order to estimate the oscillation frequency we estimate the power spectrum density (PSD) associated to the time evolution of the mean membrane potential *v* for heterogeneous (Δ_*J*_ = 0.02) and homogeneous (Δ_*J*_ = 0) case subject to noise of the same amplitude, namely *σ* = 0.00842.

For the heterogenous case, we observe that the two neural mass models and the network simulations agree quite well among them as evident in Fig. 19. The only noticeable difference are the positions of the main peak of the PSD, that are slightly different. The main peak of the 3rd order neural mass model and of the network simulations are located both around *f*_0_ ≈ 50.79 Hz, while the 2nd order neural mass reveals a peak located at *f*_0_ ≈ 50.95 Hz, as visible in the inset of Fig. 19.

**Figure 19.**
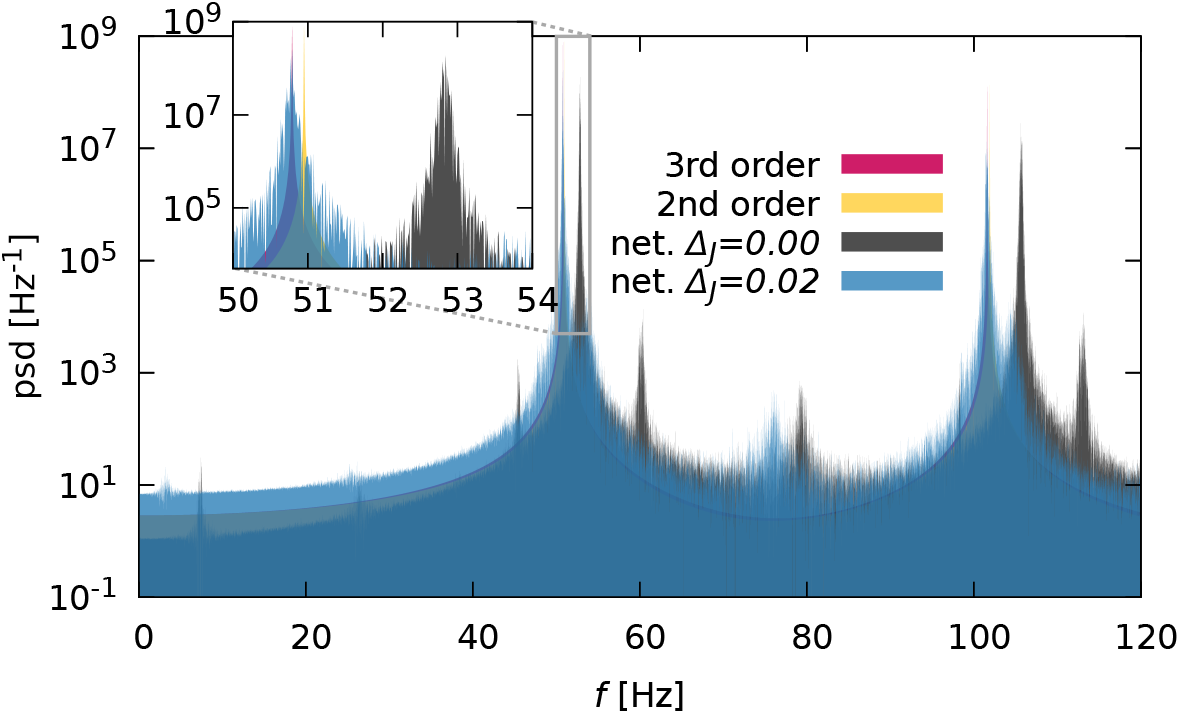
Power spectrum density (PSD) versus the the frequency *f* of the mean membrane potential *v* for the 3rd order neural mass model (red), the 2nd order neural mass model (orange) and the network simulation of size *N* = 200000 with Δ_*J*_ = 0 (black) and Δ_*J*_ = 0.02 (blue), respectively. The colors are plotted with transparency to allow the visualization of overlapping parts. The inset shows a zoom around the global maxima. All the other parameters are fixed as in Fig. 1 and the noise amplitude to *σ* = 0.00842. The PSD are measured via a discrete Fourier transform of a time series measured over 100 seconds with a step size of 2.5 × 10^−3^ ms.

The simulations of the network with homogeneous couplings result in a PSD with a peak at a higher frequency *f*_0_ ≈ 52.8 Hz, still in the vicinity of the heterogenous peaks.

The PSD around the peak is a bit broader for the network simulations compared to the neural mass results, since the network presents also finite-size fluctuations. In the network simulations, homogeneous and heterogeneous, we observe also peaks at combinations of the first two harmonics, not present in the neural mass models, suggesting that finite-size effects can lead to combinations of these harmonics similar to the beating phenomenon.

By analyzing the microscopic dynamics of the network for the same parameters in the homogenous case, we observe that the histogram *n*_*ν*_ of the single neuron firing rate *ν*_*i*_ is extremely localized with a peak around *ν*_0_ ≈ 19.27 Hz (*ν*_0_ ≈ 26.41 Hz) for the asynchronous (clustered) state (see Fig. 20). These data confirm that in the clustered regime the neurons mostly fire every two population bursts, since the frequency of COs is *f*_0_ ≈ 52.8 Hz. In the heterogenous case, the situation is more complex, as shown in Fig. 20 (b) the histogram of the firing rates has a main peak at *f*_0_*/*2 with symmetric tails at lower and higher frequencies and a secondary peak at *f*_0_, where *f*_0_ ≈ 50.79 Hz. These data confirm the previously reported analysis for the heterogeneous model performed in Subsection 4.1.4. The heterogeneity in the couplings is essentially responsible for the distributed firing rates. Furthermore, the most part of the neurons fire every second bursts, but a small group is locked to the population activity.

**Figure 20.**
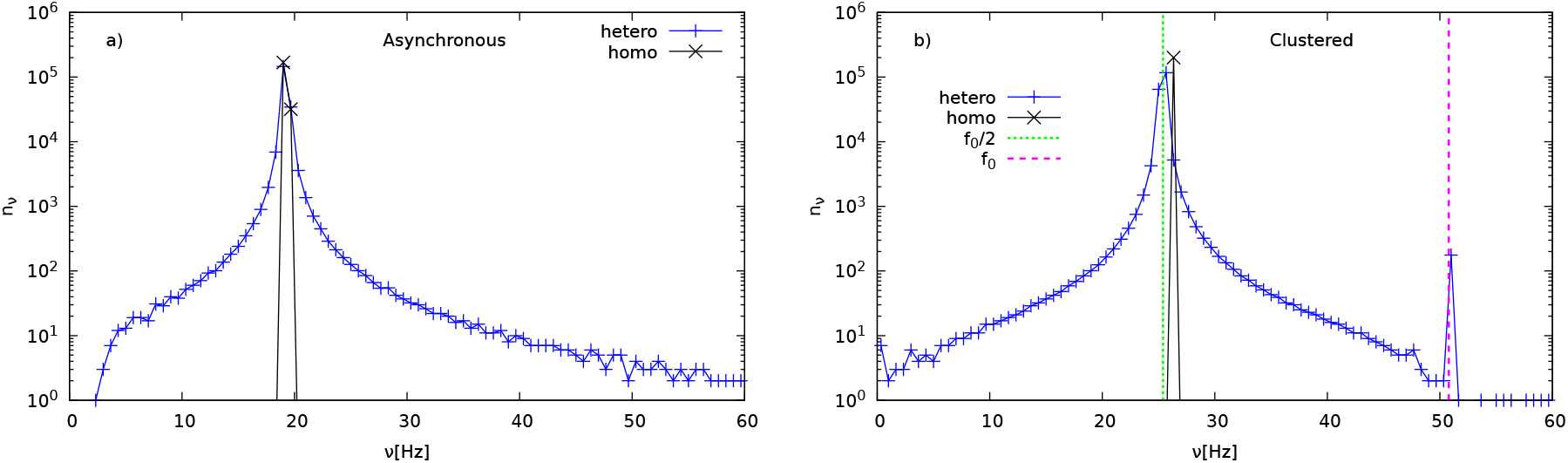
Histogram *n*_*ν*_ of the single-neuron firing rate for the homogeneous and heterogeneous networks for the asynchronous (a) and clustered (b) phases. Black (red) line refers to homogeneous (heterogenous) networks. All parameters as in Fig. 19.

Increasing the noise amplitude only changes the frequency slightly, e.g., a noise of *σ* = 0.03 results in a frequency around 52.44 Hz for the 3rd order mean-field neural mass, this means that we observe *γ*-oscillations in the whole region of coexistence.

## 5. Summary and outlook

We have shown that for a globally coupled inhibitory network of QIF neurons the presence of independent Gaussian noise promotes the emergence of COs both for heterogeneous and homogeneous synaptic couplings. The observed oscillations emerge at some critical noise amplitude *σ*_*c*_ via a sub-critical Hopf bifurcations giving rise to a region of coexistence among stationary and oscillatory dynamics. For the homogenous (heterogenous) case the coexistence is observable from zero (a finite) noise amplitude up to *σ*_*c*_.

In the heterogenous case the analysis is based on the comparison of the results obtained via a direct integration of large spiking QIF networks and of the corresponding neural mass models. In the examined noise range we observe a quite good agreement among network simulations and mean-field results obtained via the pseudo-cumulant expansion arrested to the third order [21]. The second-order neural mass model fails to reproduce the simulations for sufficiently large noise amplitudes.

The neural mass model (at the third order) captures well the macroscopic behaviour of the network induced by noise and heterogeneity, however being a mean-field model cannot reproduce the microscopic dynamics, for this we should rely on numerical simulations.

The observed collective oscillations are population bursts, where roughly half of the neurons fire in alternation in correspondence of each single collective event. However, there are irregularities to this behavior. In the heterogeneous case we observe that initially the rate at which the surviving neurons emit a spike in an irregular manner decreases over time and successively it becomes constant, i.e., it becomes a Poissonian process in the long run. The origin of the initial behaviour is due to the heterogeneity in the synaptic couplings, while the following phase is due to the presence of the Gaussian noise. Indeed in the homogenous case we observe only the second phase.

Furthermore, for heterogenous couplings we observe that the regular behaviour of the neurons, i.e. firing every two population bursts, is observable only for synaptic couplings corresponding to the median of the distribution. For sufficiently large inhibitory couplings we have silent neurons, while neurons displaying a 1:1 locking with the population bursts are observable in a wide interval of synaptic couplings.

In order to characterize the stability of the asynchronous regime we have introduced a new criterion based on the long-term evolution of the system, once the stationary configuration corresponding to the asynchronous regime has been subject to a non-infinitesimal global deformation. This criterion allows to identify the basins of attraction of the two coexisting regimes, therefore resembling the basin stability analysis [24]. Furthermore, the method captures with very good accuracy the Hopf and saddle-node bifurcation points delimiting the coexistence regime in the heterogenous and homogenous cases. This criterion can represent a useful alternative to the linear stability analysis and find application in the context of complex networks for the characterization of their dynamical regimes.

The nature of the noise is fundamental in order to observe the reported phenomena. Indeed for Lorentzian distributed white noise it was shown that the corresponding low-dimensional neural mass model [43, 44] exhibits only a stable foci and no oscillatory regime, which we have also verified via network simulations.

Clustering phenomena similar to the one here analysed have been reported in [5] for globally coupled inhibitory homogenous networks for conductance based and current based neural models in presence of Gaussian noise. However, at variance with our model the authors considered post-synaptic potentials of finite duration and not instantaneous synapses, as in the present case. The emergence of COs in absence of a delay or of a finite synaptic time scale is peculiar of inhibitory QIF networks, as previously shown in [47].

As we have shown for the chosen parameters the frequency of the COs is in the *γ*-band, therefore the present model can be employed to analyze the emergence of transitory *γ*-bursts coexisting with asynchronous dynamics observed in many experiments [14, 15, 16, 17]. In particular, our model can represent a more realistic alternative to the damped harmonic oscillator driven by noise employed in [16] to reproduce the emergence of spontaneous *γ*-cycles in awake primate visual cortex (V1). Finally, the indicators we have introduced in Paragraph 2.3.3 to characterize the regularity/irregularity of the single-neuron dynamics with respect to the global activity can find applications in the analysis of spiking events with respect to the Local Field Potential evolution in experimental data.

## Acknowledgments

We thank D.S. Goldobin for extremely useful interactions and N. La Miciotta, S. Olmi, A. Pikovsky for valuable discussions as well as Yvonne Feld for helping us in realizing several figures. The simulations were performed at the HPC Cluster CARL, located at the University of Oldenburg (Germany) and funded by the DFG through its Major Research Instrumentation Program (INST 184/157-1 FUGG) and the Ministry of Science and Culture (MWK) of the Lower Saxony State. This work also used the Scientific Compute Cluster at GWDG, the joint data center of Max Planck Society for the Advancement of Science (MPG) and University of Gottingen. Y.F. received financial and logistical support by the German Academic Scholarship Foundation (Studienstiftung des Deutschen Volkes). A.T. received financial support by the Labex MME-DII (Grant No. ANR-11-LBX-0023-01), by the ANR Project ERMUNDY (together with Y.F.) (Grant No. ANR-18-CE37-0014), and by CY Generations (Grant No ANR-21-EXES-0008), all part of the French program Investissements d’Avenir. Y.F. thanks the Istituto dei Sistemi Complessi (CNR) in Sesto Fiorentino, Italy and the Laboratoire de Physique Théorique et Modélisation, CY Cergy Paris Université, in Cergy-Pontoise, France for the hospitality offered during 2022 and 2023, where part of this work has been developed.

## Appendix A. Implementation of the numerical simulations

For the heterogenous case, in order to compare the results of the network simulations with the neural mass models [21] we consider synaptic couplings following a LD *h*(*J*_*i*_) with median *J*_0_ and half width at half maximum (HWHM) Δ_*J*_, that we fix deterministically as follows

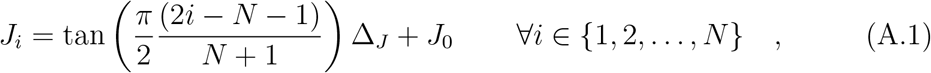

to avoid spurious effects related to extreme values of the couplings and to allow for a faster convergence at sufficiently large system sizes towards the corresponding mean-field results as previously shown e.g. in [20, 48]. It should be noticed that even for a LD centered at a quite negative value, namely *J*_0_ = −20 as in our case, a small number of positive coupling is expected. For the parameter values used in this paper (Δ_*J*_ = 0.02) the percentage of excitatory synaptic coupling is 0.032 %, therefore we can consider their influence on the macroscopic dynamics as negligible.

The initial values of the membrane potentials are deterministically chosen as in Eq. (23) from the LD expected in the thermodynamic limit (6) for the asynchronous state. Note that, to avoid correlations between *V*_*i*_ and *J*_*i*_ that would result from the previous deterministic equations, the list of *J*_*i*_ values is shuffled before creating the initial state of the network.

In order to analyze the transition from the asynchronous to the partially synchronized regime due to the noise, we perform simulations where the noise amplitude is varied quasi-adiabatically. In particular, we start from some initial noise value, typically *σ* = 0, and we simulate the models for a certain time interval *t*_*S*_, after discarding a transient time *t*_*T*_ . The quantities of interest are evaluated only during the interval *t*_*S*_. Then we increase the noise amplitude by an amount Δ*σ* and we repeat the previous procedure by an initial condition that is the last configuration obtained at the previous step. The noise is increased in steps of amplitude Δ*σ* up to some maximal value is reached. Then the procedure is repeated by decreasing the noise at each simulation step by Δ*σ* until the initial noise value is recovered.

As already mentioned, for the QIF model the threshold value would be *V*_th_ = +∞ and the reset one *V*_re_ = −∞, it is possible to take in account exactly the integration among these extrema in absence of noise if the neurons are supra-threshold by employing event driven techniques [49, 50]. However, in presence of noise we should perform usual clock driven simulations by employing finite threshold and reset values as suggested in [20].

In particular, we implement the finite threshold crossing and the spike emission as follows. Whenever *V*_*i*_(*t*) *> V*_th_, the neuron enter in a refractory period of duration *T*_*R*_ = 2*/V*_*i*_, after this phase the membrane potential is resetted to *V*_*i*_(*t* + *T*_*R*_) = −*V*_*i*_(*t*). Thus employing a variable resetting value related to the neuron evolution, this avoids spurious synchronization phenomena induced by using the same reset value for all neurons as suggested in [20]. Furthermore, the neuron *i* will fire at a time *t* + *T*_*R*_*/*2 that approximately corresponds to the time it would reach +∞ as shown in [20]. Somehow, the usage of finite thresholds and reset value is less matematically accurate, but it reflects more the dynamics of real neurons [51].

To simulate the network model we have numerically integrated the stochastic differential equation Eq. (1) by employing a clock driven scheme. In particular we have employed the Heun method [52] for its higher accuracy in the treatment of the determistic part with respect to a standard Euler scheme.

The iterative Heun method [52] applied to our network model reads as :

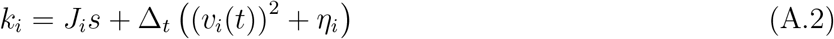

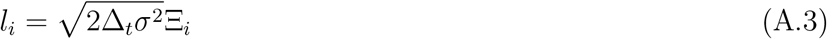

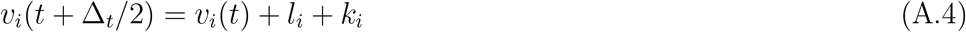

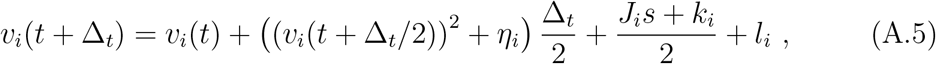

where *V*_*i*_(*t*) are the membrane potentials, *k* and *l* are auxiliary variables, and Ξ_*i*_ is a Gaussian random number with zero mean and unitary standard deviation that is drawn separately for each neuron. Morever Δ_*t*_ is the integration time step, whose choice will be discussed in the next Appendix, and *s* represents the network activity and it is the number of spikes emitted in the network in the interval Δ_*t*_ divided by *N* .

## Appendix B. Selection of the integration time step

For the numerical integration of the network model we need to select an optimal time step Δ_*t*_, which should lead to high accuracy in the integration joined to a minimal computational cost.

In deterministic systems this choice is quite simple, one just select the largest time step for which the integrated orbits converge to the same value up to some accuracy. In a stochastic system this cannot happen, therefore we rely on a different concept.

In the present case, we know from the mean-field approach that in the asynchronous regime for sufficiently small noise values *σ < σ*_*SN*_ the system should always relax towards a PDF of the membrane potential that is a LD, namely

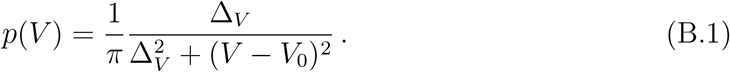

with *V*_0_ = *v*^*^ and Δ_*V*_ = *πr*^*^, where (*v*^*^, *r*^*^) are the fixed point solutions of the neural mass model Eq. (16). Therefore, we considered a small noise value *σ* = 0.001 *< σ*_*SN*_ and for different integration time steps Δ_*t*_ we have verified if the distribution of the membrane potentials converge to (B.1) or not.

In particular, we initialized the simulation always with membrane potetials distributed as in (B.1), then we simulate the system for a time interval of 1.2 secs and every 0.12 ms we accumulate the instantaneous values of the membrane potentials in a histogram of 5000 bins with *V* ∈ [−100, 100]. We do not consider the neurons in their refractory periods to prevent from unphysical overestimations of large *V* values. From the final histogram we obtain the PDFs shown in Fig. B1 for three different Δ_*t*_*/τ*_*m*_ = 1 × 10^−3^; 3 × 10^−3^; 6 × 10^−3^ together with the expected PDF (B.1).

As evident from Fig. B1, the larger time-step leads to clear artifacts in the estimation of the PDF. Already by considering Δ_*t*_*/τ*_*m*_ = 3 × 10^−3^ leads to a noticeable improvement, in particular the right inset of Fig. B1 reporting *p*(*V*) in linear scale around the maximum show essentially no differences among (B.1) and the estimated PDFs with Δ_*t*_*/τ*_*m*_ ≤ 3 × 10^−3^. However, in the semi-logarithmic scale (left inset and main figure) the numerically estimated PDF for Δ_*t*_*/τ*_*m*_ = 3 × 10^−3^ still presents numerical artifacts for sufficiently negative *V* values.

For a time step Δ_*t*_*/τ*_*m*_ = 1 × 10^−3^, we cannot notice any artifact and essentially we have a perfect coincidence with the theoretical PDF (B.1). We can already conclude that this time step give a sufficient accuracy to the simulations, however to be on the safe side we opted for Δ_*t*_*/τ*_*m*_ = 2.5 × 10^−4^.

**Figure B1.**
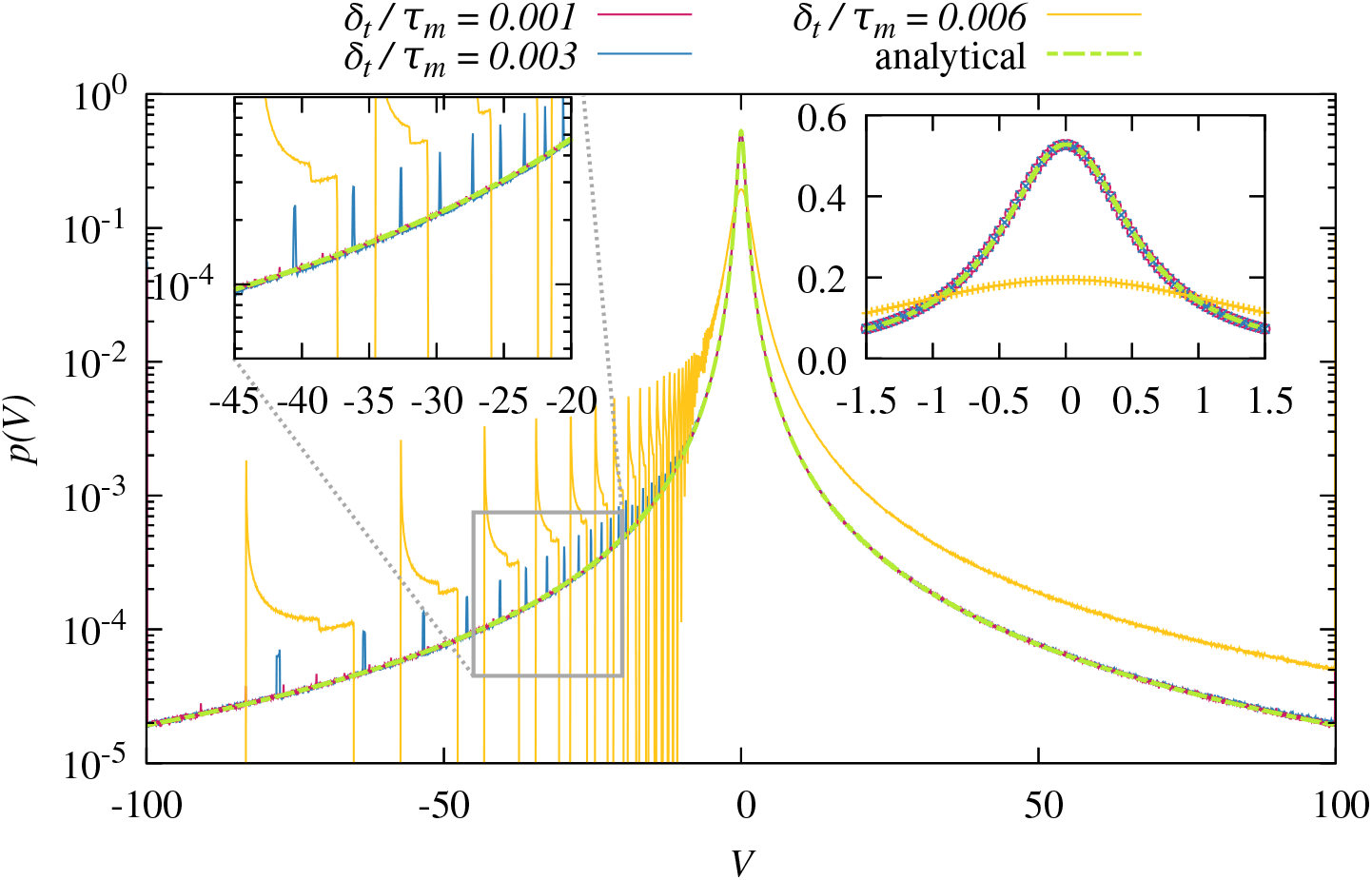
Probability density function *P* (*v*) as measured for a network of size *N* = 200000, Δ*J* = 0.02 and *σ* = 0.001. The dashed lime-green line shows the analytical prediction while the solid lines represent the measurements for a step-size of 0.001 (red), 0.003 (blue) and 0.006 (yellow). The left inset shows a zoom, while the right inset shows a range around the peak in linear scale and uses symbols for each data point that are color-coded as above.

